# Structural asymmetry in biotic interactions as a tool to understand and predict ecological persistence

**DOI:** 10.1101/2023.01.25.525558

**Authors:** Alfonso Allen-Perkins, David García-Callejas, Ignasi Bartomeus, Oscar Godoy

## Abstract

A universal feature of ecological systems is that species do not interact with others with the same sign and strength. Yet, the consequences of this asymmetry in biotic interactions for the short- and long-term persistence of individual species and entire communities remains unclear. Here, we develop a set of metrics to evaluate how asymmetric interactions among species translate to asymmetries in their individual vulnerability to extinction under changing environmental conditions. These metrics, which solve previous limitations of how to independently quantify the size from the shape of the so-called feasibility domain, provide rigorous advances to understand simultaneously why some species and communities present more opportunities to persist than others. We further demonstrate that our shape-related metrics are useful to predict short-term changes in species’ relative abundances during seven years in a Mediterranean grassland. Our approach is designed to be applied to any ecological system regardless of the number of species and type of interactions. With it, we show that is possible to obtain both mechanistic and predictive information on ecological persistence for individual species and entire communities, paving the way for a stronger integration of theoretical and empirical research.

## Introduction

Ecologists seek to obtain a mechanistic understanding of the processes governing the dynamics of ecological systems. The meaning of *mechanistic*, however, varies across ecological levels of organization. For instance, while biotic interactions are key for a mechanistic understanding of community dynamics (Levine *et al*., 2017), so is plant growth for population ecologists or photosynthesis rates for ecophysiologists (Peñuelas *et al*., 2011). Nevertheless, a common trade-off across fields is that obtaining such mechanistic understanding often comes at the cost of losing predictive ability (Dietze, 2017; Petchey *et al*., 2015; Tredennick *et al*., 2021). These mismatches between understanding ecological processes and predicting ecological dynamics are due to multiple causes, yet a common limitation is the lack of correspondence between the mathematical tools used for understanding and those used for predicting. For instance, in community ecology, while the mechanistic understanding of complex dynamics of interacting species uses well-established population models (Bimler *et al*., 2018; Godoy & Levine, 2014), predictive studies are much less developed, and generally use a completely different set of tools including machine learning techniques (Civantos-Gómez *et al*., 2021; Evans *et al*., 2011).

In order to break the methodological trade-off between understanding and predicting, recent approaches have applied tools from the structuralist approach (Saavedra *et al*., 2020; Song *et al*., 2020). In mathematics, structural stability is a property of dynamical systems that defines the range of conditions compatible with stable dynamics. Applied to ecological communities, the structuralist approach posits that the structure of species interactions influences the opportunities for multiple species to persist in the long term and, therefore, their coexistence (Godoy *et al*., 2018; Saavedra *et al*., 2017). The larger these opportunities (measured as the size of the so-called feasibility domain), the larger the range of environmental conditions the community can withstand without losing any species. Among other insights, it has been predicted (Saavedra *et al*., 2017), and empirically shown (Bartomeus *et al*., 2021; García-Callejas *et al*., 2023) that this situation occurs when species experience strong self-regulation and niche differentiation. These insights are probabilistic in nature (Cenci *et al*., 2018), coherent with the principle that is impossible to quantify the whole set of abiotic and biotic factors modulating community dynamics (Shoemaker *et al*., 2019).

The occurrence, sign, and strength of biotic interactions are arguably the most relevant information needed to understand and predict ecological outcomes at the species and the community level (Macarthur, 1957). As such, biotic interactions are the raw material that the structuralist approach uses to define the feasibility domain, which in turn allows predicting the persistence of whole communities (i.e., community feasibility), species composition, or abundances at equilibrium (Tabi *et al*., 2020). For instance, it has been shown that in both random and empirical communities, the size of the feasibility domain (i.e., the range of species growth rate values compatible with persistent communities) is dependent on the type of interaction, mean interaction strength, and connectance (Grilli *et al*., 2017). However, the distribution of interaction strengths is not only relevant to determine the size, but equally important, the shape of the feasibility domain. Conceptually speaking, if all species interact in the same way the feasibility domain will be symmetrical, but in practice, interactions among species in real-world systems are predominantly asymmetric (Adler *et al*., 2018; Bascompte *et al*., 2006). That is, species generally differ in their pairwise per-capita effects *α*_*ij*_ ≠ *α*_*ji*_. These asymmetries in interactions translate to asymmetries in the feasibility domain, as recently exemplified in communities of different types (Grilli *et al*., 2017) (see an additional example in Suppl. Section 1).

Understanding the asymmetry of the feasibility domain is important because it implies that not all species have the same opportunities to coexist. That is, some species are more affected than others by random changes in environmental conditions, such as interannual variations in precipitation. Therefore, even in communities with large feasibility domains, it is likely that particular species may become locally extinct when environmental conditions change. These more vulnerable species represent the “weak spot” for the feasibility of the whole community (Grilli *et al*., 2017). This process was exemplified in a recent study by Tabi *et al*. (2020) who, working with protist communities in temperature-controlled environments, showed that in communities with more asymmetric feasibility domains, it is harder to predict which protist species went extinct first. Overall, these examples have shown that it is not only important to understand the potential of the community structure to maintain all of its constituent species, but also the relative vulnerability of each species to perturbations in their demographic performance.

Despite these recent studies providing solid and rigorous progress, it is still poorly understood how to independently quantify the shape and the size of the feasibility domain. This separation is critical to better predict ecological outcomes at the level of species and communities. Conceptually, we should aim for a shape measure that is independent of the size of the feasibility domain. Although these properties are not fully independent (see an example in Suppl. Section 2), it is currently not explored to what extent they can be meaningfully disentangled and interpreted. For instance, when measuring the shape of the feasibility domain as the heterogeneity in the distribution of the side lengths of its borders (Grilli *et al*., 2017), we are implicitly measuring the change in the size of the domain as well. This is because a mild increase in the variability of the side lengths of the feasibility domain implies also a change in its size. A similar conceptual problem applies to characterizing the asymmetry of the feasibility domain by computing the variability of direct effects between species pairs (Tabi *et al*., 2020). To advance in our ability to predict the dynamics of ecological communities, here we provide new developments that control for the correlation between the shape and size of the feasibility domain. Failing to control for this relationship may significantly hinder our ability to estimate species’ vulnerabilities to exclusion in local communities.

Coupled with these methodological advances, we require empirical tests to evaluate their usefulness. Otherwise, they will remain valid theoretical constructs but with limited application to real conditions. Predictions of ecological dynamics using the structuralist approach have focused so far on either computer simulations (Grilli *et al*., 2017) or highly controlled laboratory (Tabi *et al*., 2020) and experimental settings (Bartomeus *et al*., 2021), yet hard tests under field conditions are scarce (Song & Saavedra, 2018). These tests are key to shed light on the applicability of the perspective we present here to predict species dynamics across a wide range of ecological systems and environmental conditions. The litmus test of ecological prediction is that of predicting species abundances (Clark *et al*., 2020; Tredennick *et al*., 2017), which is relatively uncommon due to its inherent complexity, as it encompasses a greater degree of information than species richness or composition (McGill *et al*., 2007).

In this perspective, we develop and test new probabilistic metrics to understand species’ relative vulnerability to exclusion from local communities, and the relationship of these metrics with temporal changes in observed abundances. In particular, we build from recent developments that quantified the minimum random perturbation that can exclude a species from a feasible community (denoted as “full resistance” in Medeiros *et al*. (2021) and as “robustness” in Lepori *et al*. (2023)). We propose to call this metric “exclusion distance”, which is a definition rooted in the analysis of exclusion probabilities that follows. We expand on this idea and develop a novel theory that quantifies: 1) the absolute and relative probabilities of being the first species excluded from the community (referred to as “probability of exclusion” and “exclusion ratio”, respectively), and 2) an intuitive index that quantifies the asymmetry in species’ vulnerabilities to extinction in any ecological community (referred to as the “asymmetry index”), regardless of their overall feasibility potential. Importantly, this set of metrics can be applied to ecological communities with arbitrary interaction types and number of species. Lastly, we ask whether these metrics can inform about changes in species population growth rates through time in a grassland community composed of a diverse set of species with contrasted functional roles. We show that the metrics developed here are phenomenologically correlated with year-to-year variation in observed population growth rates across species.

## Theoretical framework

### Preliminaries: Definition of the feasibility domain and its size

To understand the theoretical background of this perspective, we introduce some key concepts of the structuralist approach. This framework posits that the structure of species interactions determines the opportunities for species to coexist by conditioning the size of the feasibility domain. Particular structures of species interactions, such as intraspecific competition exceeding interspecific competition, promote larger sizes of the feasibility domains and, therefore, larger opportunities for species to coexist (Barabás *et al*., 2016; García-Callejas *et al*., 2023). In order to measure the size of the feasibility domain, we need to connect the structure of species interactions with a model describing the dynamics of interacting species. The underlying model commonly used is the linear Lotka-Volterra (LV) model, as it represents a balance between tractability and complexity, and can generate population dynamics with the range of complexity of more complex models (Cenci & Saavedra, 2018; Hart *et al*., 2018; Saavedra *et al*., 2017). On the one hand, the model allows summarizing the effect of the abiotic environment on the performance (i.e., the balance between mortality and resource intake) of individual species into a single parameter, the intrinsic growth rate (Saavedra *et al*., 2020). On the other hand, despite its simplicity, the LV model has successfully explained and predicted the dynamics of diverse ecological systems under controlled and natural experimental conditions (García-Callejas *et al*., 2021; Godoy & Levine, 2014; Saavedra *et al*., 2020; Tabi *et al*., 2020; Venturelli *et al*., 2018). The model is of the form:

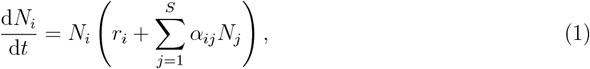

where *N*_*i*_ denotes the abundance of species *i, S* is the number of species, *r*_*i*_ is the intrinsic growth rate of species *i* which represents how species *i* grows in isolation under given abiotic conditions, and *α*_*ij*_ is the element (*i, j*) of the interaction matrix **A** and represents the per-capita biotic effect of species *j* on the per-capita growth rate of species *i*.

In a given ecological community, the necessary conditions to sustain long-term positive abundances for all species depend on its interaction matrix **A**, and are known as the feasibility domain of the community, henceforth FD. Formally, the FD is the parameter space of intrinsic growth rates that leads to a feasible (positive) equilibrium, i.e., the set of **r** such that **N\*** = −**A**^−1^**r** *>* **0** for non-singular interaction matrices **A** (i.e., for matrices whose inverse, denoted as **A**^−1^, exists), where **r** is a column vector whose *i*-th element is the intrinsic growth rate *r*_*i*_, and **N**^*^ is a column vector whose *i*-th element represents the abundance of species *i* when the feasible equilibrium exists, 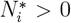 (Song *et al*., 2018). When the points in the FD are also dynamically stable (i.e., when the real part of the eigenvalues of the linearized Jacobian matrix of Eq. (1) is negative), then feasibility ensures the stable coexistence of the *S* species (Logofet, 2005). Despite real-world communities might no be dynamically stable, especially when in communities with high interaction asymmetries (Bunin, 2017), it is possible to find feasible systems with unstable equilibria. In such systems, it should be possible to evaluate at least the short-term population dynamics in response to temporal reconfigurations of the structure of biotic interactions (Medeiros *et al*., 2023). This is the particular case of annual communities (García-Callejas *et al*., 2021), which is the system we empirically test the metrics presented here.

According to the structuralist approach, the larger the size of the FD, the more likely the community can persist without any species going extinct. To quantify its size, previous work has assumed no a-priori knowledge about how the intrinsic growth rates will change within a community. For that reason, we assume that all parameter values are equally likely to be observed in the entire parameter space (Saavedra *et al*., 2020), and use the probability of uniformly sampling a vector of feasible intrinsic growth rates on a representation of that space, the unit-ball *B*^*S*^. In mathematical terms, the probability that all species can persist, Ω(**A**), is equal to the ratio of the following volumes:

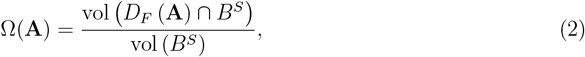

where *X* ∩ *Y* denotes the intersection of any two sets X and Y (Gourion & Seeger, 2010; Saavedra *et al*., 2020, 2016a; Song *et al*., 2018). Hereafter, we denote the relative volume Ω as FD *size*. To illustrate these mathematical expressions for a community with three species, Fig. 1 shows the intersection of the FD with the unit-ball. Specifically, only the points that lie within those triangles represent feasible vectors of growth rates. The vertices contained in Fig. 1 (colored circles) depict vectors of intrinsic growth rates where only a single species has positive equilibrium abundances 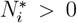 (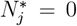 for *i* ≠ *j*), whereas each edge shows vectors of growth rates where a given species (the species of the vertex with the same color as the edge) goes excluded. Importantly, both communities in Fig. 1 have FDs with the same size Ω = 0.064 but differ in their shape. For communities with *S* species, the size Ω(**A**) lies between 0 and 0.5 (Song *et al*., 2018). Thus, the closer Ω(**A**) is to 0.5, the larger the likelihood that the structure of species interactions **A** promotes a feasible community regardless of the differences in intrinsic growth rates among species. The size Ω(**A**) presented in Eq. (2) can be efficiently computed as follows:

**Fig. 1:**
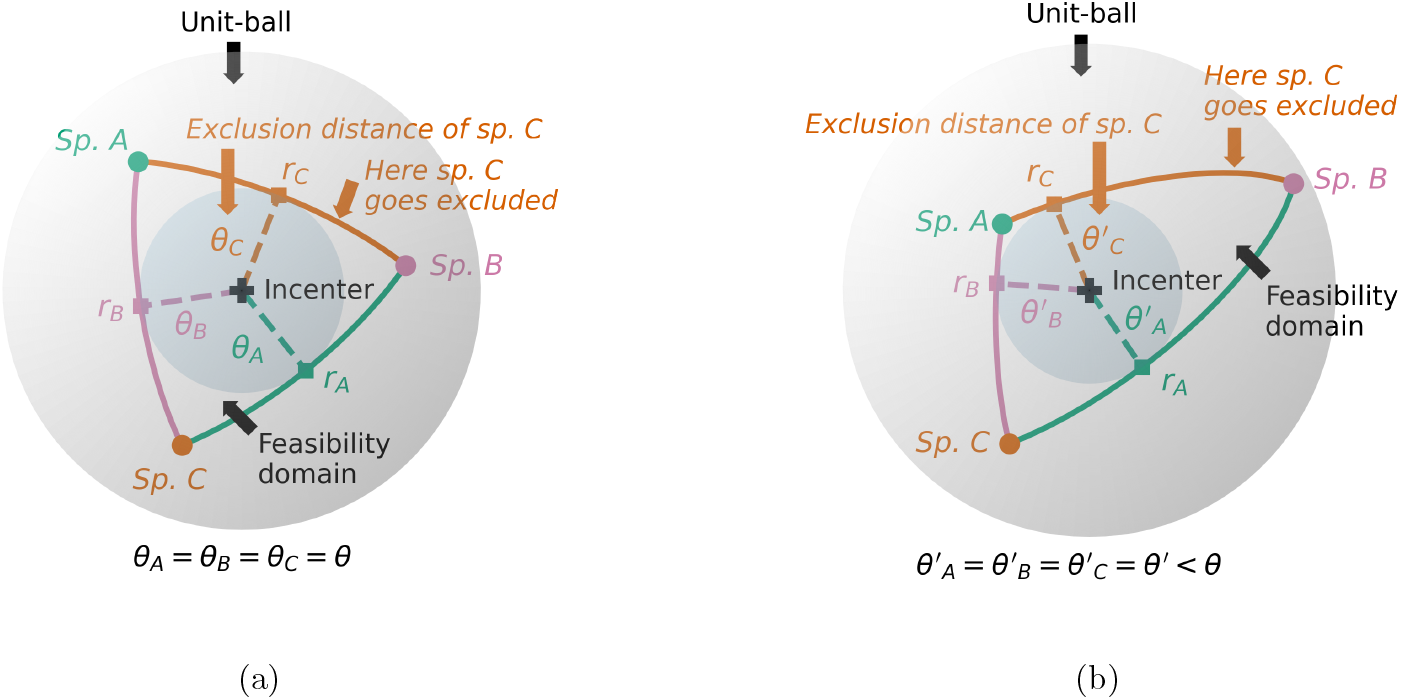
Examples of feasibility domains (FDs) for communities with *S* = 3 species. The white sphere represents the unit-ball, whereas the spherical triangles show the borders of the FD. Within the spherical triangles lie the vectors of intrinsic growth rates that lead to the persistence of the three species altogether. In this example, the size of both triangles (Ω) is the same (Ω = 0.064). However, in panel (a) we represent a symmetric domain (equilateral spherical triangle), whereas panel (b) shows an asymmetric one (scalene spherical triangle). The distance between the incenters of the FD (black plus markers) and their respective borders (square markers) are represented with dashed colored lines. In addition, the blue spherical caps are a guide to the eye to compare that the minimum distance from the incenter to the edge of the FD is shorter in the asymmetric triangle. The distance between the incenter and the border is *θ*_symmetric_ = 0.417 in panel (a), and *θ*_asymmetric_ = 0.365 in panel (b).

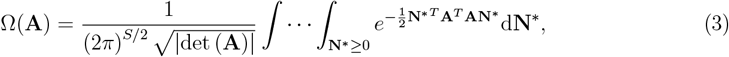

via a quasi-Monte Carlo method (Genz & Bretz, 2009; Saavedra *et al*., 2016b; Song *et al*., 2018).

### Feasibility domain’ shape: Interpreting perturbations as distances

The FD’s size Ω can be used as an indicator of the tolerance of a community to random environmental variations. However, two communities with the same number of species *S* and identical values of Ω can still have very different responses to perturbations of their intrinsic growth rates **r**, depending on the shape of their FDs (Fig. 1). Because this shape is critical to understand and predict the feasibility of the whole community and the vulnerability of its constituent species, we here build on one recently proposed metric (the species exclusion distance (Lepori *et al*., 2023; Medeiros *et al*., 2021)) to develop three novel metrics that thoroughly characterize such shape, namely: 1) the species’ probability of exclusion, 2) their exclusion ratios and, at the community level, 3) the asymmetry index (see Box 1 for a summary of these metrics).

#### Box 1

Metrics based on the shape of the feasibility domain.

***Species exclusion distance***. Given a feasible community whose dynamics can be well approximated with a linear LV or models that can be summarized via a LV structure (Eq. (1) and Godoy & Levine (2014); Saavedra *et al*. (2017)), the exclusion distance of species *i*, denoted as *θ*_*i*_, represents the minimum perturbation of the community’s initial vector of growth rates (**r**_0_) that can exclude species *i* from that community (Lepori *et al*., 2023; Medeiros *et al*., 2021). The geometrical definitions and reasoning required for its analytical calculation (Eq. (S7)) are presented in Suppl. Sections 3 and 4.

***Probability of exclusion***. This probabilistic metric, denoted as 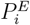, ranges between 0 and 1, and estimates the average (absolute) probability that species *i* will be the first one to be excluded if a destabilizing perturbation took place in a feasible community under LV dynamics. According to its formal definition (Eqs. (4) and (6)), on the one hand, the calculation of these probabilities does not require an initial vector of growth rates. On the other hand, their values mainly depend on the shape of the FD. In ideal symmetric communities with *S* species, all species are equally likely to be the first species excluded and, consequently, the probability of exclusion of each species is given by 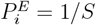.

***Species exclusion ratio***. Given a community with *S* species, the exclusion ratio of species *i*, denoted as *ER*_*i*_, is a relative metric that assesses whether the probability of exclusion for species 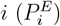 is large by comparing it to that of an ideal symmetric community with the same number of species (1*/S*) (Eq. (5)). By definition, this metric ranges between 0 and infinity, and is equal to 1 only in symmetric communities. The larger the exclusion ratio of species *i*, the larger the relative probability that *i* can be the first species that is excluded.

***Asymmetry index***. This community-level metric describes the distribution of the aver-age probabilities of exclusion 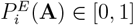 and, thus, the (a)symmetry of the FD’ shape. To quantify it, we use a commonly-used equitability index (sensu Tuomisto (2012)): the relative Shannon diversity index of the average probabilities of exclusion 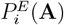(**A**) (Eq. (7)).

To introduce these metrics, we start by answering a simple question: What is the maximum perturbation in environmental conditions a community can withstand without losing species and therefore becoming unfeasible? To answer this question, first, we should consider that species go excluded when the vector of growth rates of the community **r** moves from its initial position until it reaches the border of the FD. The further away a vector of intrinsic growth rates **r** is from the border of the FD, the larger the perturbations of **r** the community can withstand while being feasible. The largest minimum distance between the growth vector **r** and the border occurs when the vector **r** is at the *incenter* of the FD ℑ (black plus markers in Fig. 1) because its minimum distance to any edge of the FD is the same regardless of the direction of the perturbation (i.e isotropic perturbation). Thus, the maximum isotropic perturbation that a community can cope with, while being feasible, will be geometrically equivalent to the distance from the incenter to all the edges of the FD (along the surface of the unit-ball), denoted by *θ* (dashed lines in Fig. 1).

Now we can ask how the shape of the FD influences the maximum perturbation of the incenter ℑ. In general, when two communities follow Eq. 1 and have the same number of species *S* and equal sizes of their FDs, the more asymmetric a FD is, the closer the edges are to the incenter ℑ and, consequently, the smaller the minimum distance between them *θ* (i.e., the smaller maximum perturbation). This difference is illustrated in our example for *S* = 3 species in Fig. 1, where the maximum isotropic perturbation from the incenter is larger for the symmetric domain than for the asymmetric one (*θ*_symmetric_ = 0.417 and *θ*_asymmetric_ = 0.365, respectively). The calculations required to find the incenter ℑ and the maximum isotropic perturbation *θ* of a given interaction matrix **A** can be found in Suppl. Section 3. Nevertheless, comparing the FD of communities from empirical studies is not straightforward, as communities may vary both in species richness and in the degree of interaction asymmetry between species. This variation in the size and number of species will consequently affect the shape of the FD. The next sections are therefore dedicated to explaining under realistic conditions how changes in the shape of the FD allow us to understand and predict the magnitude of perturbations that species can withstand (exclusion distance) and the probability of exclusion for species and entire communities.

### Feasibility domain’ shape: Species exclusion distances

In the previous section, to answer the question of which is the maximum random perturbation a community can handle without losing species, we considered the distance from a specific point, the incenter (I), to the edges of the FD (*θ*). However, the same question can be explored by taking any other initial vector of growth rates within the FD **r**_0_ ≠ ℑ. In this more general case, we can find the point in the border of the FD (**r**_0,min_) at a minimum distance to the initial vector of growth rates (**r**_0_). That distance between **r**_0_ and the closest point in the border **r**_0,min_, *θ*_0_, defines the maximum perturbation that will cause one of the *S* species to be excluded. The advantage here is that the approach can further inform us about which species will go extinct first: The identity of such species only depends on the location of **r**_0,min_ on the border of the FD. Following the example in Fig. 2(a), the closest point to the initial vector of growth rates **r**_0_ (blue circle) that is located in the border of the FD is **r**_*C*_ ≡ **r**_0,min_ (orange square), and it is on the edge where species *C* is excluded (orange edge).

**Fig. 2:**
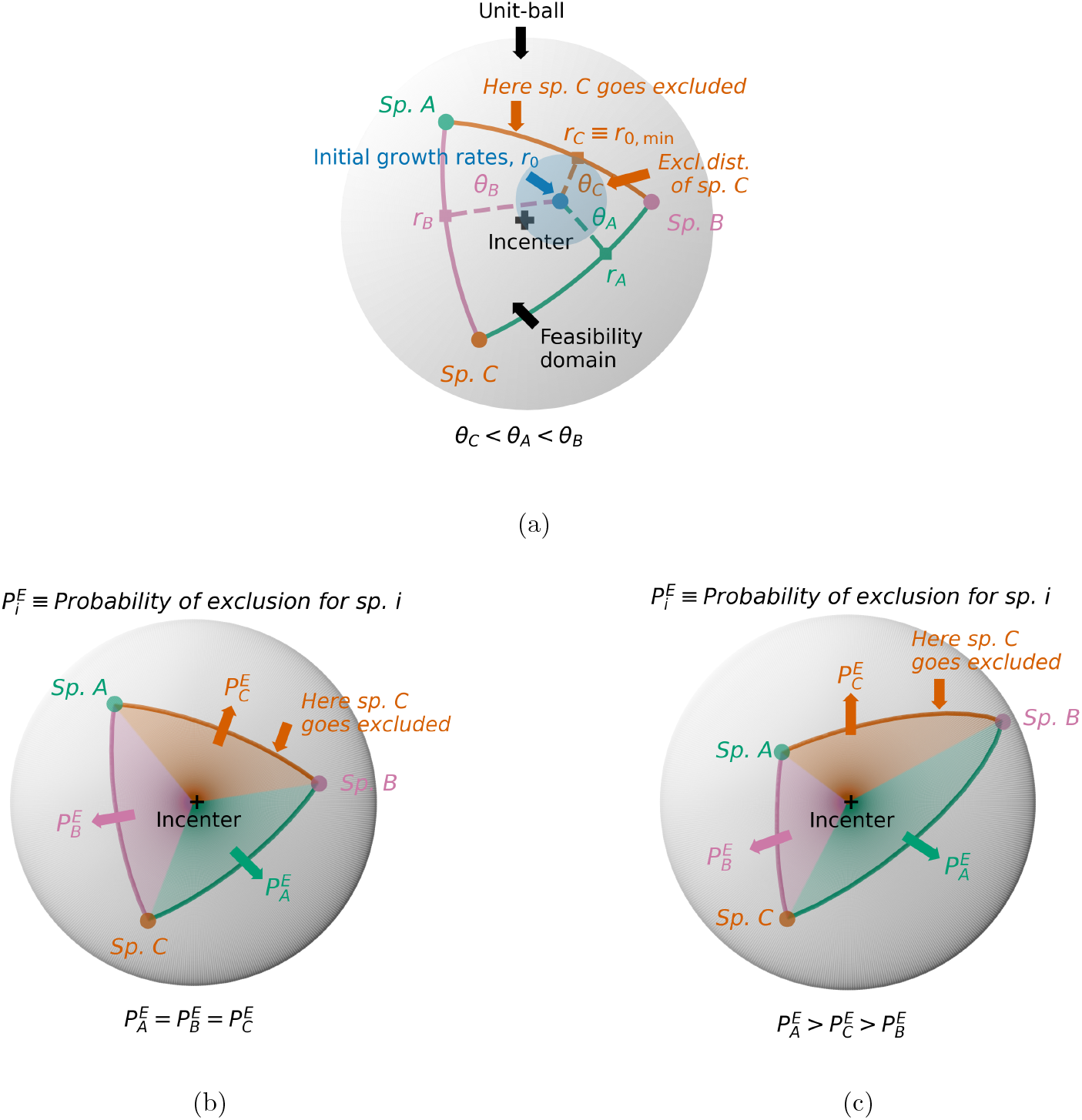
(a) Exclusion distances *θ*_*i*_ and maximum perturbation *θ*_0_(≡ *θ*_*C*_) for the community in Fig. 1(a) when the initial vector of growth rates **r**_0_ (dark blue dot) is not at the incenter (black plus marker). Dashed lines represent the exclusion distances (i.e., the shortest distances along the unit-ball from the initial vector of growth rates (*r*_0_) to the border where a given species goes extinct (square markers)). The colors of squares and dashed lines are those of the species excluded. The exclusion distances depicted in this example are *θ*_*A*_ = 0.407, *θ*_*B*_ = 0.546, and *θ*_*C*_ = 0.239, respectively. (b) Representation of the average probabilities of exclusion for the species in the FD shown in Fig. 1(a): 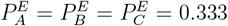. The colors correspond to those of the species that will be the first ones to be excluded. (c) This panel shows the same information in Fig. 2(b) for the FD in Fig. 1(b). In this asymmetric FD, the average probabilities of exclusion are 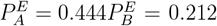, and 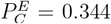, respectively. The size of all the FDs included in this figure is the same (Ω = 0.064).

In many situations, such as in conservation and restoration efforts, we are interested in the persistence of particular species, which poses the question of whether we can calculate such maximum perturbation before losing a given species *i* from the community. To do so, we simply need to calculate the minimum distance (along the unit-ball) from **r**_0_ to the edge of the FD where *i* goes excluded, denoted by *θ*_0,*i*_. This distance is what we define as *species exclusion distances*. In Fig. 2(a), we represent the exclusion distances for the initial vector of growth rates **r**_0_ with dashed lines.

### Feasibility domain’ shape: Probabilities of exclusion and exclusion ratios

In order to estimate the species exclusion distances *θ*_*i*_ we need quite a lot of information on the system, namely: the structure (matrix) of species interaction **A**, which defines the FD, along with a vector of intrinsic growth rates, **r**_0_. However, characterizing such vector can be a challenging, labor-intensive work even in small communities (Bartomeus *et al*., 2021), and, hence, this information is not available for many communities, usually those including animal and long-lived species. In these cases, we can use the above definitions and a probabilistic perspective to estimate 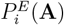, the probability that species *i* will be the first one to be excluded if a strong perturbation took place:

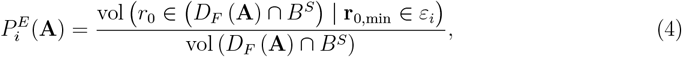

where *ε*_*i*_ represents the edge of the FD in which species *i* goes excluded. Note that this probability is formulated independently from the precise specification of initial conditions, similar to the structural forecasting in Saavedra *et al*. (2020). According to Eq. (4), the probability 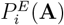 corresponds to the proportion of the FD’s volume where the vectors of intrinsic growth rates are closer to the edge where species *i* is excluded (Fig. 2). Therefore, by definition, 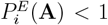 for *S >* 1 (when **A** is non-degenerate). We name the quantity 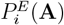 as the *probability of exclusion for species i*.

The probability of exclusion of a given species depends on the size and shape of the FD. As Fig. 2 illustrates, such probabilities are mainly conditioned by the distance between the species vertices (circles) and the incenter (plus markers): the closer the vertex of a given species is to the incenter ℑ (compared to other vertices), the larger the probability that such species will be the first one to be excluded, and therefore the whole community will not be longer feasible. This prediction is important because it implicitly indicates that the shape of the FD can be characterized by computing the average probabilities of exclusion. For instance, in communities where all species interact in the same way, and consequently have symmetric FDs, all vertices are at the same distance from the incenter, and the probability of exclusion is the same for all species 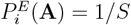 (see Fig. 2(b) for an example). In contrast, the larger the variation across the probabilities of exclusion, the larger the asymmetry of the FD because species present also asymmetries in their interaction matrix (e.g. effect on species *i* on *j* is not the same as the effect on species *j* on *i*). Our approach allows to assess whether the probability of exclusion for a given species is larger compared to a situation in which all species interact in the same way and therefore are equally likely to be the first species excluded:

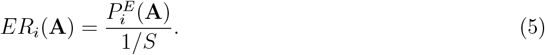

We denote such quantity as the *species exclusion ratio* (García-Callejas *et al*., 2023) (see Suppl. Section 5 for a generalization of Eq. (5)). If *ER*_*i*_(**A**) *<* 1 (*ER*_*i*_(**A**) *>* 1), then the probability of exclusion of species *i* is smaller (larger) than that of a symmetric community. So we hypothesize that the smaller the value of *ER*_*i*_(**A**), the larger the species probability of persistence.

Finally, to compute the average probabilities of exclusion 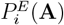 in Eq. (4) and the exclusion ratios derived from them in Eqs. (5) and (S9), we propose the following approach:

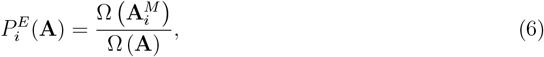

where 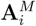 represents a modified interaction matrix, obtained by replacing the vertex of species (*V*_*i*_) by the incenter, ℑ, which is equivalent to replace the *i*−th column of **A** by the negative of the incenter’s column vector. Intuitively, the solid angle 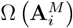 is the proportion of the volume of the unit-ball where the vectors of growth rates are closer to the edge where species *i* is excluded (i.e., **r**_0,min_ ∈ *ε*_*i*_. If we consider the examples in Figs. 2(b) and (c), we simply say that the area occupied by a given color (e.g. green) is equal to the area obtained when placing the green vertex of species *A* in the position of the incenter.

It is worth noting that by using the same quasi-Monte Carlo method applied to estimate Eq. (3), we can compute efficiently both the numerator and denominator in Eq. (6), and the results obtained are in excellent agreement with those obtained by simulating the dynamics of the underlying LV model (Eq. 1) (see Suppl. Section 6 for details on the LV simulations, as well as examples in communities with three and more species). This might seem a technical detail but it has important implications for the application of our metrics by a broad range of ecologists. The problem we solve here is that finding several initial conditions randomly for estimating 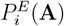 from numerical simulations is a time-intensive task when the size of the FD is small (Ω ≲ 10^−12^) with orders of magnitude that vary from minutes-hours with our approach to days-weeks with the numerical LV estimation (Fig. S7). On top of this problem, solving the LV model numerically can be far from trivial when the abundance of certain species grows indefinitely, as in communities with mutualistic interactions and weak self-regulation. For those reasons, the estimation of the average probabilities of exclusion 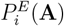 from Eq. (6) (via quasi-Monte Carlo method) is –to the best of our knowledge– the most efficient alternative to computing species-level probabilities of exclusion from a structuralist point of view.

### Feasibility domain’ shape: Asymmetry index

Taking into account the features of the probabilities of exclusion 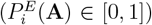 and their link with the shape of the FD, we use a relative Shannon diversity index *J* ^*′*^, introduced by Pielou (1976), to describe the distribution of the average probabilities of exclusion. This index, which in our context we called “asymmetry index”, is a commonly-used equitability index in ecological studies (Tuomisto, 2012). It has the following three desirable properties: i) it is maximal and equal to one when the average probabilities of exclusion are equal (i.e, when the FD is symmetric); ii) it approaches zero when the average probabilities of exclusion are very dissimilar; and iii) the index shows values in the middle of its scale when the differences between probabilities can be intuitively considered “intermediate” (see a full description of its features in (Smith & Wilson, 1996)). The asymmetry index is of the form:

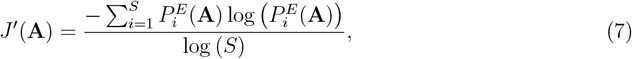

for a community with *S* species. We use the notation *J* ^*′*^(**A**) to highlight the dependence of the results on the interaction matrix. Since 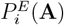 depends on the size of the FD, the latter also affects *J* ^*′*^(**A**). Indeed, given FDs whose sides are proportional, the larger the size of a given domain, the greater the differences among 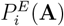 and, thus, the more asymmetric (i.e., the smaller *J* ^*′*^(**A**)) (Fig. S8 for an example). Nevertheless, the dependence of 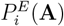 and *J* ^*′*^(**A**) on size is smaller than that of other shape indices. In particular, we tested the relative change in shape indices in situations in which the size of the FD changes while its shape is kept approximately constant (Suppl. Section 7 for details). We show that, for communities of different sizes, the asymmetry index *J* ^*′*^(**A**) is much less sensitive to variations in size than two other indices, namely the variance of the cosine of the side lengths as developed by Grilli *et al*. (2017), and the variance of the niche differences developed by Saavedra *et al*. (2017) (Fig. 3).

**Fig. 3:**
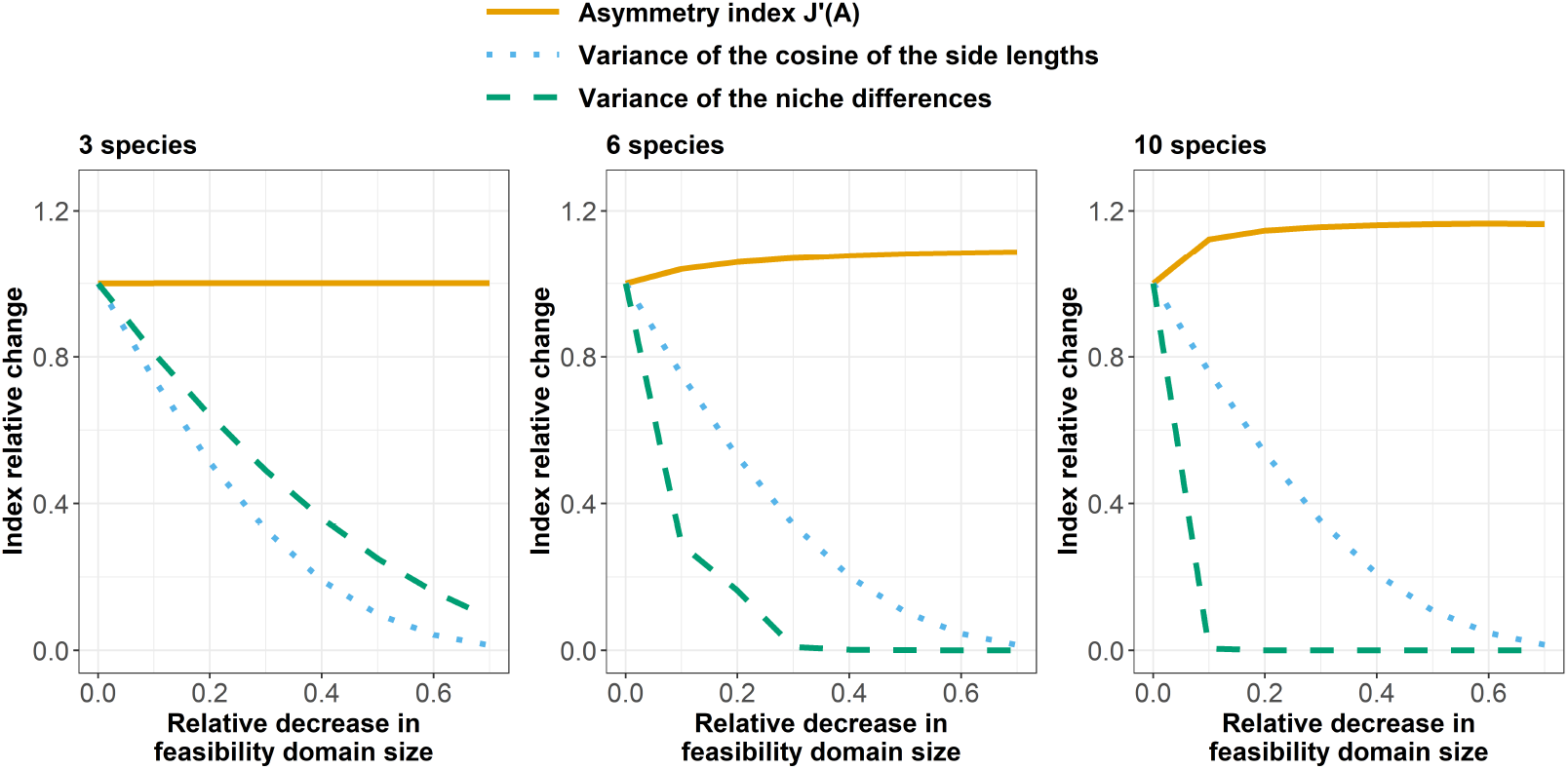
Dependence of several shape-indices on the size of the feasibility domain (FD) and on species richness. For three, six, and ten species we show how much the value of the three indexes evaluated vary (y-axis) when we reduce the size of the FD but preserve the proportions of the distances between the domain’s vertices (x-axis). The indexes shown are: the asymmetry index *J* ^*′*^(**A**) (continuous lines), the variance of the cosine of the side lengths developed by Grilli *et al*. (2017) (dotted lines), and the variance of the niche differences developed by Saavedra *et al*. (2017) (dashed lines). A value of index relative change equal to one means that the index value is maintained across reductions of the FD. The metric we present in this perspective (“asymmetry index”) slightly varies as we account for more species in the community but this variation is negligible compared to previous indices available in the literature. A graphic example of how the size of the FDs is reduced while maintaining the proportions among its sides is given in Suppl. Section 7.

### Applications to empirical communities

With the perspective presented above and the proposed structural metrics, we can advance our understanding of the feasibility, composition and short-term dynamics of ecological communities. Importantly, this approach is valid for interactions among species belonging to a single trophic level, i.e. communities driven by competition and facilitation, as well as for communities with relevant interactions across trophic levels, such as food webs or bipartite networks. We provide in the following sections two complementary analyses of natural communities. First, we expand on the use of our approach to predict short-term population growth rates using data from a highly-resolved annual plant community. Second, in Box 2, we evaluate the extinction risk of plants and pollinators based on the structure of their bipartite interactions.

### Predicting plant population dynamics

Our main prediction is that the asymmetry of species interactions, which can be computed with the above-described shape-based metrics, modulates the population dynamics of species, and therefore, can inform of temporal changes in species abundances and population growth rates, at least in the short-term. In addition, it can also inform on the population growth rate expected variability across time. If these hypotheses hold, our approach can help predict changes in abundances in species-rich communities while maintaining a mechanistic understanding (Ehrlén & Morris, 2015). In particular, our first theoretical expectation is that, the smaller the exclusion ratio (*ER*_*i*_) or the larger the exclusion distance (*θ*_*i*_) of a given species, the larger the perturbation needed to exclude it from the community. Hence, it is more likely that these species will exhibit positive short-term growth rates because they will tend to increase their populations (Fig. 4). Complementing this hypothesis, we also expect that those species with smaller exclusion ratio (*ER*_*i*_) will present higher variation in population growth rates, while this variation will be reduced in species with larger exclusion ratio (*ER*_*i*_) (Suppl. Section 8). This is because, for species presenting a small *ER*_*i*_, the perturbation can occur in many different directions, either promoting or hindering population growth across time, but is less likely to occur in the specific direction that promotes the extinction of such species. Conversely, for a species presenting a large *ER*_*i*_, any perturbation is more likely to head in the multiple directions that promote its exclusion, so in order to persist, such species with large *ER*_*i*_ will only withstand small perturbations. This implies small changes in population growth rates in the long-term (Arnoldi *et al*., 2018). Finally, we predict that the more accurate the structural characterization of the community (by providing information on species interactions and their intrinsic growth rates), the better the prediction. That is, we expect that predictions based on detailed structural information, such as the exclusion distances *θ*_*i*_ (extracted from species’ intrinsic growth rates at a given time), will perform better than those derived from a probabilistic average, like the exclusion ratios *ER*_*i*_ (Fig. 4).

**Fig. 4:**
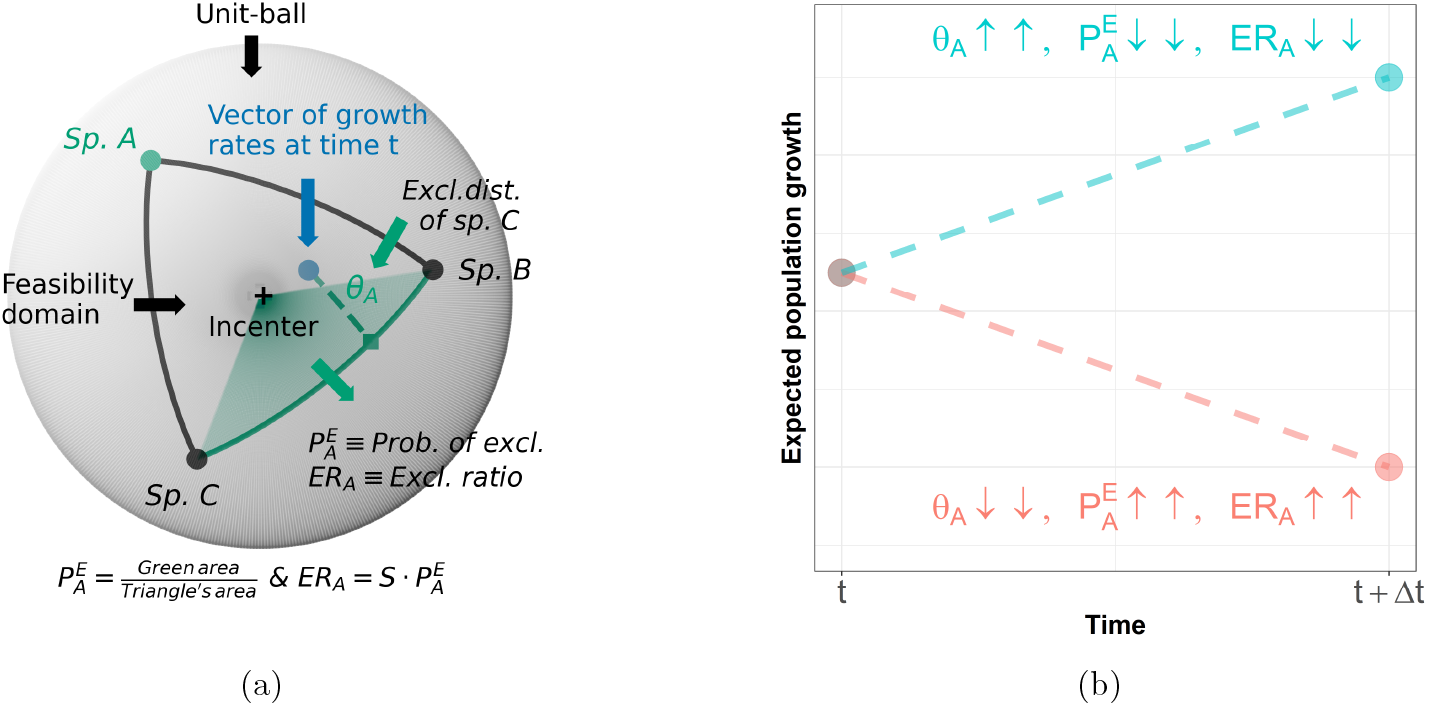
(a) Example of the shape-based exclusion metrics that can be estimated for a species that belongs to a (symmetric) community with *S* = 3 species, namely: the exclusion distance of species *A* (*θ*_*A*_), its probability of exclusion 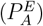 and its corresponding exclusion ratio (*ER*_*A*_). (b) Expected relationships between the shape-based exclusion metrics of a single species in a given year (*t*) and its future population growth (*t* + Δ*t*).

Empirical information to test our three central predictions comes from a detailed observational study in species-rich plant communities from a Mediterranean grassland. There, we gathered information on the observed abundances, intrinsic reproduction rates, and interactions between annual plant species during seven years, from 2015 to 2021 (García-Callejas *et al*., 2021).

### Study system

From November 2014 to September 2021, we sampled each year at Caracoles Ranch, located in Doñana National Park, SW Spain (37 °04’01.0” N, 6 °19’16.2” W), 19 annual species naturally occurring in the study area, that accounted for *>* 90% of plant biomass in the community. These 19 species belong to disparate taxonomic families and exhibit contrasted functional profiles along the growing season (Supplementary table S1). The earliest species with small size and open flowers, such as *Chamaemelum fuscatum* (Asteraceae), peak at the beginning of the growing season (February), while late species with succulent leaves, such as *Salsola soda* (Amaranthaceae), grow during summer and peak at the end of the growing season (September-October).

In order to parameterize empirical estimates of intra- and interspecific pairwise plant-plant interactions (i.e., the elements *α*_*ij*_ of the interaction matrices **A**) we related species’ individual reproductive success to the number of potentially competing individuals of each species within plant neighborhoods for each year, which allow us to extract the seed production in the absence of neighbors *λ*, the growth vector **r** and **A** (see further details (García-Callejas *et al*., 2021; Lanuza *et al*., 2018), and in Suppl. Section 10). Aside from these data, we also recorded annually an independent measure of field abundance for each of the 19 species considered (Suppl. table S1). The sampling coverage of plant-plant interactions, quantified via rarefaction curves, was *>* 99% in all annual communities (Suppl. Section 12).

### Estimation of shape-based metrics

Using the empirical estimates of the interaction matrices and intrinsic growth rates, we obtained the annual exclusion distances and the exclusion ratios for the 19 plant species in our system. The former was derived from Eq. (S7), whereas the latter was estimated from Eqs. (3) and (5), via quasi-Monte Carlo methods based on Eq. (6), and generating 10,000 random draws for each required value of exclusion probability, 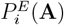. For these calculations, we assumed that the interaction coefficients and the exclusion metrics represent the corresponding averaged values within a given year, while these average values vary across years due mainly to inter-annual variation in rainfall. In addition to the structural metrics, we tested whether the interaction matrices of our system were diagonal-stable (Suppl. Section 11). Our analyses showed no evidence of diagonal stability. Hence, we cannot ensure that dynamical stability is fulfilled in addition to feasibility. This apparent contradiction does not affect the following analyses, because we explored year-to-year changes in population growth rate and its relationship with exclusion metrics based on interannual realizations of the interaction matrix. Although long-term dynamical stability might not be relevant for our understanding of the short-term dynamics of annual plant communities in which their biotic interactions are reset each hydrological year, it is important to bear in mind that proving such dynamical stability is a necessary condition for predicting which species will persist at long-term equilibrium Saavedra *et al*. (2017).

### Statistical analyses

We analyzed the relationship between the observed annual growth rate in our grassland study system, defined as *N*_*i,t*+1_*/N*_*i,t*_, and two of our metrics: the species exclusion ratio *ER*_*i*_ (based only on the interaction structure) and the species exclusion distance *θ*_*i*_ (based on both the interaction structure and the demographic performance). Note that abundance data is completely independent of the data used to calculate the interaction strengths (see details in (García-Callejas *et al*., 2021)). Therefore, they provide a rigorous test to evaluate the ability of our metrics to predict the population dynamics of interacting species.

We developed two sets of regression models to test the relationship of annual population growth rate with our metrics. For the first set of six models, we implemented Generalized Linear Models (GLMs) with Gamma and Gaussian distributions with log-link function, and took as single explanatory variables, respectively, the number of individuals of species *i* in the previous year, and two shape metrics of species *i* from the community in the previous year: the exclusion ratio *ER*_*i*_(**A**_*t*_) and the exclusion distance *θ*_*i*_(**A**_*t*_). The second set of models is conceptually similar, but we fitted a negative exponential relationship (Non-lineal least squares fit, NLS), in practice by using log_10_ (*N*_*i,t*+1_*/N*_*i,t*_ + 1) as the response variable, following visual observations of the scatter-

plot. We further tested for the inclusion of temporal autocorrelation structure that allows for unequally spaced time observations (CorAR1 in R-package `nlme` v3.1-152 Pinheiro *et al*. (2021)), as well as species identity as a random effect. Then, we compared those models on the basis of AIC to asses which model better described the patterns observed (Burnham & Anderson, 2004). For these analyses, given that not all the estimated interaction matrices contained the same pool of plant species, we only used the results for those species that were present from 2015 to 2021. In addition, we did not consider the data of the growing season of 2018 (that is, we did not try to predict the population growth in 2018 from the data in 2017) because that growing season was dominated by an extreme flooding event, which does not reflect the annual plant dynamics (see García-Callejas *et al*. (2021) for details on that perturbation and its effects on the community). After applying the above filtering, we selected seven plant species from contrasted taxonomic groups, namely: *Beta macrocarpa, Centaurium tenuiflorum, Hordeum marinum, Leontodon maroccanus, Parapholis incurva, Polypogon maritimus*, and *Salsola soda*.

Overall, in these models we assumed that the relevant abiotic and biotic factors that drove the species dynamics during a given year were encoded in our shape-based metrics (Ehrlén & Morris, 2015). In addition, note that our models did not include information on the dispersal ability of the focal species, but given the nature of the study system which flooded every year, we assume there is no limitation to global dispersal in the study area.

Our analyses were conducted in R, with the `stats` v4.1.0 (R Core Team, 2021) and `lme4` v1.1-28 (Bates *et al*., 2015) packages. We also checked model assumptions with the R-package `nlstools` v2.0-0 (Baty *et al*., 2015).

## Results

In our seven years of sampling, we documented and parameterized 126 and 2,142 intra- and interspecific pairwise plant-plant interactions, respectively. From them, we extracted 7 different annual interaction matrices **A**, whose average number of species per year was 15.29 *±* 3.35. The estimated interaction strengths *α*_*ij*_ (the elements of **A**) were either negative (i.e. denoting competition) or positive (i.e. denoting facilitation). Further, the distribution of the absolute value of *α*_*ij*_ was right-skewed throughout all the years, with a majority of comparatively weak interactions (Suppl. Fig. S11). The variation of those asymmetries in interaction strengths, as expected, also exerted changes in the size and shape of the communities’ FD. This result is consistent with the idea that species sensitivity to perturbations is dependent on the system state and its structure of interactions with other species, as in the case of food webs (Beauchesne *et al*., 2021) or systems out-of-equilibrium (Medeiros *et al*., 2023). The maximum values of FD size (Ω) and symmetry (*J* ^*′*^) were reached from 2016 to 2018 (Ω_max_ = 0.382 and 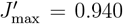), and the minimum ones in 2019 and 2020 (Ω_min_ = 0.0823 and 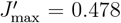) being 0.5 and 1 the upper bounds of Ω and *J* ^*′*^ (Suppl. Fig. S12). In general, the vulnerability to exclusion of the seven species present every sampled year, expressed via their exclusion ratios and exclusion distances, fluctuates across years (Suppl. Figs. S13, S14, S15).

Results from our models supported our theoretical expectations that our shaped-based metrics allow describing short-term population growth dynamics and their variance across time. Models that included shape metrics provided a better fit than the model only including abundance from the previous year, suggesting that the metrics considered carry a predictive signal. In particular, a negative-exponential least squares fit better described the observed relationships, as confirmed by comparing AIC values of the different models (table S13). Therefore, we subsequently present the results obtained from the exponential NLS fits. These models show that the higher the exclusion ratio, the lower the growth rate of the following year (effect-size: -0.31, table S9). That is, those species that have larger probabilities of exclusion reduced their population growth rate the following year. Conversely, the exclusion distances have a positive effect on population growth rate (effect-size: 3.33, table S12) indicating that those species that were further away from the edge of the FD increased their population growth rate the following year. We hypothesized that metrics with more detailed structural information would be better predictors of population growth rate, i.e., models including exclusion distances would provide better fits than models based on exclusion ratios or raw probabilities of exclusion. This was not the case, as models incorporating different metrics showed similar AIC values, within a range of 63.862 *±* 0.6192 units (table S13). Here we therefore focus on the model with exclusion ratio as the predictor, which showed the lowest AIC values and allows for a clear ecological interpretation (Fig. 5). In that model, analyzing the observed and predicted population growth allows a qualitative categorization of the observations in three groups. Those years in which species presented an exclusion ratio between 0 and 0.75 showed positive population growth rates on average and a greater variance, as predicted by our second hypothesis and supported by our simulations (Suppl. Fig. S10). Conversely, we observed flat or negative population growth rates as well as lower variance for species that showed an exclusion ratio *>* 0.75 in a certain year, particularly if the exclusion ratio was *>* 1.25. This suggests that, in our system, an exclusion ratio of 0.75 qualitatively differentiates species showing contrasted population growth trends across time. Lastly, accounting for species as random intercepts or for temporal autocorrelation did not improve model fits.

**Fig. 5:**
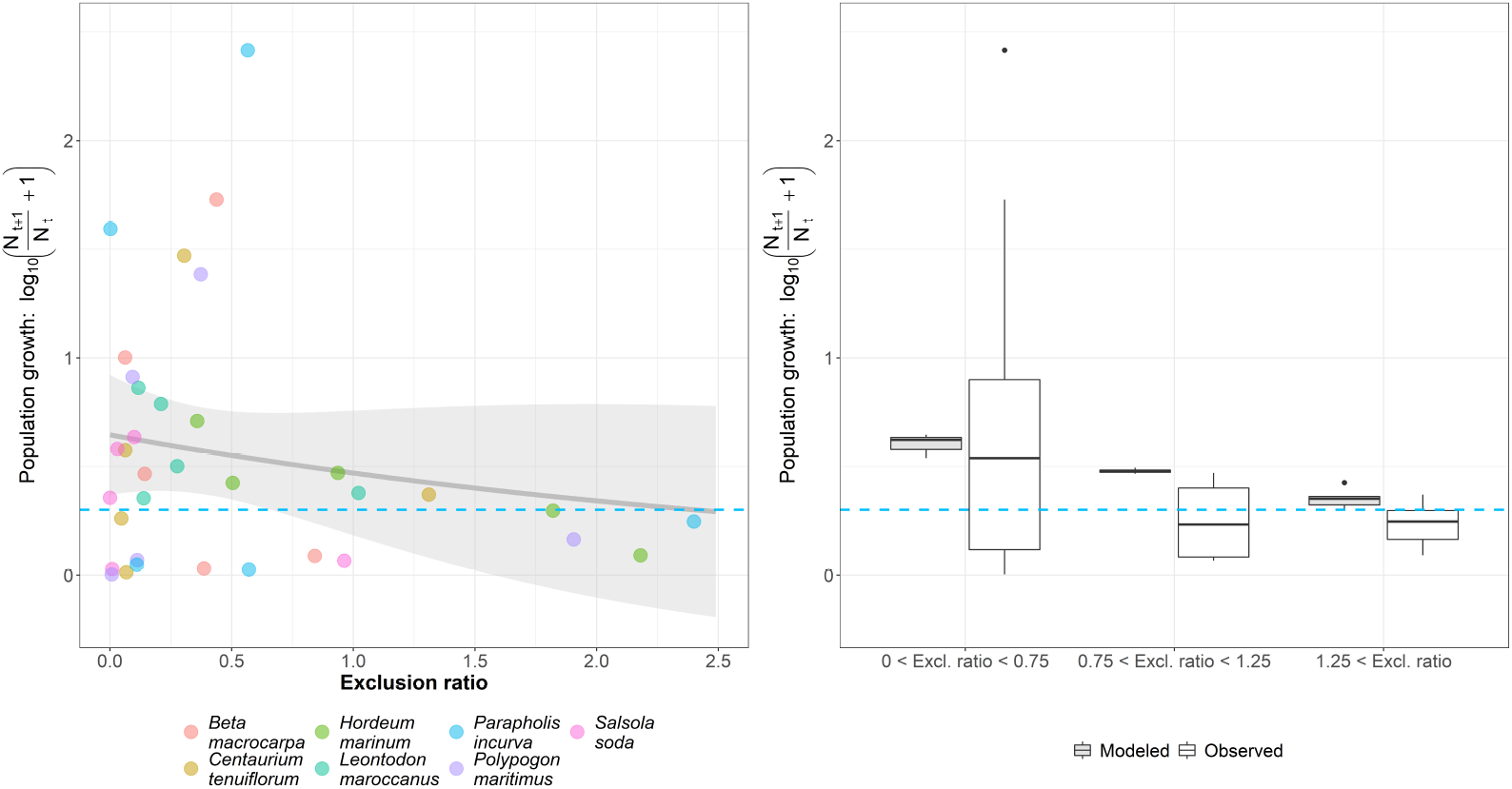
Results for the non-linear model log_10_ (Pop. growth + 1) = *a ·* exp (*b ·* Excl. ratio), showing the effect of the exclusion ratio in a given year *t* on observed population growth rate *N*_*i,t*+1_*/N*_*i,t*_. The left panel shows the observed values of exclusion ratio and population growth for each studied plant species (colored dots), and the values predicted by the NLS model (grey line), and their 95% confidence interval (grey shaded area). Above the blue dashed line population growth increases (*N*_*i,t*+1_ *> N*_*i,t*_), whereas below that line it decreases (*N*_*i,t*+1_ *< N*_*i,t*_). Box-plots in the right panel display the observed and predicted population growth across three different groups of exclusion ratios: i) *ER*_*i*_ *<* 0.75, ii) 0.75 *< ER*_*i*_ *<* 1.25 and iii) 1.25 *< ER*_*i*_.

### Box 2

Assessing exclusion risks in mutualistic communities with shape-based metrics: A proof of concept.

The structure of plant-pollinator interactions can encapsulate information about the sensitivity of different species to perturbations. For example, there is considerable evidence supporting the relationship between specialization and extinction risk (Bartomeus *et al*., 2013; Biella *et al*., 2020; Burkle *et al*., 2013; Memmott *et al*., 2007; Scheper *et al*., 2014): extinction risk declines with increasing numbers of mutualist partners, from least-linked (most specialized) species to most-linked (most generalized) ones.

Here we test if the ranking of our shape-based metrics, which quantify the species’ average probability of being the first species excluded from a given community, aligns with the expected relationship between specialization and extinction risk described in the literature. To do so, we extracted from the Mangal database (Poisot *et al*., 2016) 88 empirical plant-pollinator networks from 5 different studies with wide geographical coverage (table S14). Such data comprises 1,734 species in total, where 1,198 species were animals. On average each network had 19.70 *±* 9.18 nodes of which 13.61 *±* 6.94 were animals. For each of these networks, we estimated the communities’ effective interaction matrices by using the mean field approximation for intraguild competition and the two inter-guild parameterizations proposed in (Saavedra *et al*., 2016b) for mutualistic communities. Then, we extracted the species’ probability of exclusion, 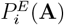 using Eq. (6). We quantified probabilities of exclusion instead of exclusion distances because we lack estimations of intrinsic growth rates in these 88 communities.

Results using the shape-based metrics show that the probability of exclusion 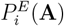 is moderately but significantly correlated with species degree (*r* = −0.37, *C*.*I*.(95%) = [−0.395, −0.338], p-value *<* 2 *·* 10^−12^) when analyzing the 88 networks together. Interestingly, our analysis identified that on average pollinators are at more immediate risk of exclusion than plants, although the trend is not statistically significant due to the large variability across in-dividual networks 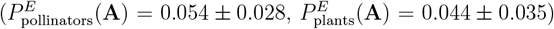. These results are in line with previous empirical and theoretical findings (Biella *et al*., 2020; Memmott *et al*., 2004). Our results were robust when considering alternative mean-field interactions among species (Suppl. Figs. S21 and S22), and are encouraging considering the potential biases and differences in sampling methodologies across networks.

## Discussion

A central theme in ecology is which information we need to mechanistically understand and predict the vulnerability of species to perturbations coming, for example, from environmental changes. This question can be approached from different angles: from looking at physiological tolerances to different environmental stressors to analyzing species’ demographic vital rates, or to exploring the joint response of species interacting with each other. In this perspective, we show that adopting a community context, supported by a reliable quantification of species interactions, is critical to predicting the effect of environmental perturbations on the persistence of both particular species and entire communities. The joint possibility of keeping a mechanistic understanding of how biodiversity is maintained across ecological levels of organization (individual species and entire communities) while achieving a reasonable predictive power is now possible by acknowledging the idea that species present in natural systems show asymmetries in their biotic interactions, and, therefore, the size and the shape of the community feasibility domain should be simultaneously explored.

A remarkable finding is that the shape-based metrics developed here are useful to predict changes in year-to-year effective growth rates among interacting species in a grassland system (see table S13). This is a strong result taking into account that the studied species are annual plants whose abundances depend on a myriad of biotic and abiotic factors including variation in precipitation, dispersion, and natural enemies (Petry *et al*., 2018). Overall, these results suggest that, although simplistic, the structure of species interactions and species’ demographic performance phenomenologically capture key ingredients of the processes controlling population dynamics and are more meaningful for making predictions that just considering abundances from previous years. Interestingly, our results suggest that experimentally deriving intrinsic growth rates, a difficult parameter to estimate empirically Bartomeus *et al*. (2021), might not be critical to obtaining reliable predictions. Further work can build on this framework to, for example, study not only the probability of exclusion but its *velocity* (Medeiros *et al*., 2021).

Overall, these results reinforce the idea that increasing system complexity is not incompatible with the predictability of the dynamics of its components (Daugaard *et al*., 2022). Rather, our results emphasize that the interaction structure of the system, which varies from year to year, implying concomitant variations in the size and shape of the FD, is in itself ecologically relevant for predicting short-term demographic performance. As shown in Box 2, this framework can accommodate more complex communities considering not only plants but also several trophic levels, such as plant-pollinator systems or multi-trophic communities (García-Callejas *et al*., 2023). Empirically documenting all relevant competitive, mutualistic and trophic interactions of a community is the next step needed to percolate the use of theoretically-informed tools to better understand and predict the temporal dynamics of ecological systems.

The metrics we propose here integrating species-level sensitivity to perturbations (i.e. the range of reproductive outcomes that lead to species persistence) within a community-level measure of feasibility (i.e. the *asymmetry index*) do not start from scratch. They rather build on previous efforts (Grilli *et al*., 2017; Medeiros *et al*., 2021; Saavedra *et al*., 2017; Tabi *et al*., 2020) that identified a key issue in the structuralist approach: the size of the FD, by itself, is not fully representative of the capacity of a community to withstand perturbations. Rather, we need to quantify its shape, i.e. the relative contribution of each species to it. Both properties are not fully independent in spherical geometry, which is the mathematical backbone of the structuralist approach (Suppl. Section 2). However, just like we might expect for two proportional triangles in two-dimensional geometry to have the same “shape” according to a quantifiable metric, we might approach that intuitive requirement in spherical geometry. Our asymmetry index fulfills this requirement more robustly than previous approaches, i.e. it’s much less sensitive to variations in the size of the FD for a given shape (Fig. 3). We provide implementations of all the metrics discussed here in the R package `anisoFun` v1.0: https://github.com/RadicalCommEcol/anisoFun, and we suggest that their adoption may help bridge the gap between ecological theory and empirical studies, in order to provide robust quantitative solutions to manage, conserve and restore biodiversity. For instance, highly asymmetric FDs due to high asymmetries in biotic interactions among species are indicative of strong differences in species probabilities of exclusion. This first-order analysis can be used as an indicator, in applied contexts, that detailed analyses of extinction risk are necessary. On the contrary, such additional risk analyses would not be so critical to perform in symmetric FDs, where all species have a similar probability of exclusion.

Both feasibility and dynamic stability are necessary conditions for the long-term coexistence of species at equilibrium (Bunin, 2017; Takeuchi, 1996). We did not find evidence of diagonal-stability in the interaction matrices of our study system, implying that although the communities are mathematically feasible, they are likely to be unstable in the long-term, i.e. external perturbations to species abundances would drive the system out of equilibrium. This notion, however, applies to long-term equilibrium analyses, which are not the main scope of our study. Rather, we are interested in short-term, year-to-year predictions of the system. In particular, our metrics can be useful in 1) predicting short-term population growths (left panel in Fig. 5) and 2) understanding its expected variability (right panel in Fig. 5). Our results show that species with high exclusion ratios have lower variability in their population growth rate across years (Suppl. Section 8), which may indicate slower recovery rates from perturbations (Arnoldi *et al*., 2018). Nevertheless, it is important to bear in mind that the exclusion distances, probabilities of exclusion, and exclusion ratios discussed here are asymptotic properties of the system, and thus remain approximations of the underlying dynamics of complex ecological communities.

The applicability of these metrics for a wide range of systems and community types remains an open question, but we can anticipate some limitations. On the theoretical side, we assume that the basic elements contained in the structure of species interactions (i.e., pairwise interactions) vary across years but can be well-approximated by annually-averaged values within a given year and that functional forms (i.e. the relationship between species abundances and overall interaction effects) are linear. However, the existence of non-linear functional forms for modeling pairwise interactions (or even the addition of higher-order interactions as in (Gibbs *et al*., 2022)) will have non-trivial consequences on system dynamics (Dougoud *et al*., 2018) and may preclude the use of the linear definition of the FD that sustains our approach. This framework is, nevertheless, able to accommodate different types of interactions beyond plant-plant interactions (Box 2), as well as alternative functional forms and dynamics (such as those of the saturating competition model by Brauer & Castillo-Chavez (2011) and other examples in (Cenci & Saavedra, 2018; Saavedra *et al*., 2017)). On the empirical side, it is premature to generalize these results to other systems because the importance of obtaining robust demographic performance data and the predictive skill gained with it is likely to be context-dependent. For instance, a single demographic performance value might not be as meaningful for systems with more complex life cycles. Overall, we argue that the toolbox developed here is ready to be used for systems whose species display similar population dynamics and similar responses to external perturbations (e.g. annual plant communities, coral reef fishes, tropical trees, plant-pollinator mutualistic systems) (Box 2, García-Callejas *et al*. (2023)).

## Conclusion

Reliable tools for predicting ecological outcomes remain elusive and, in general, disconnected from those frameworks used to understand ecological processes. Here we provide a novel set of metrics within a structuralist approach that considers the conditions leading to the persistence of species and entire ecological systems. This stems from the fact that biotic interactions among species are asymmetric in their strength and sign within ecological communities. Our overall aim with this perspective is to advance in unifying our mechanistic understanding of the maintenance of biodiversity in ecological communities while simultaneously increasing the predictive ability of their dynamics. We have shown that including information on species interactions and their demographic performance in the estimation of these shape-based structural metrics significantly improves short-term predictions of species abundances in an annual plant community, compared to predictions based only on species’ previous abundances. The species and community-level metrics presented here are general across interaction types and can be efficiently obtained for ecological communities of arbitrary richness, unlocking the possibility of systematically studying the biotic control over ecological dynamics for a wide range of levels of diversity and organisms.

## Data and code availability

The data and code used to generate the results of this study are available at https://doi.org/10.5281/zenodo.8087366 and https://github.com/RadicalCommEcol/Asymmetric_Interactions. The `anisoFun` package v1.0 is available at https://github.com/RadicalCommEcol/anisoFun and https://doi.org/10.5281/zenodo.8086505.

## Acknowledgements

We thank the Radical Community Ecology group (https://github.com/RadicalCommEcol/) for the fruitful discussions. AA-P, DGC, IB and OG were funded by the Spanish Ministry of Science and Innovation (MICINN) and the European Social fund through the MeDiNaS (RTI2018-098888-A-I00), TASTE (PID2021-127607OB-I00), and ChaSisCOMA (PID2021-122711NB-C21) projects. Additionally, OG acknowledges financial support provided by the Spanish Ministry of Economy and Competitiveness (MINECO) and by the European Social Fund through the Ramón y Cajal Program (RYC2017-23666).

## Notes

### Competing Interest Statement

The authors have declared no competing interest.

### Summary of Updates

We have performed additional simulations to better interpret our findings that there is more variability in population growth rates when species show smaller exclusion ratios. These findings allow us to better discuss the potential of the structural approach to predict short-term growth rates and their variability across species.

