## Supplementary Material for "Structural asymmetry in biotic interactions as a tool to understand and predict ecological persistence" (from another version of this preprint: version 2, 2023-05-21)

This document contains the following Supplementary Sections: "Dependence of shape on size for feasibility domains", "Calculation of the incenter and the maximum isotropic perturbation for a given feasibility domain", "Calculation of the exclusion distance of a given species", "Alternative definitions of exclusion ratios", "Estimation of the probabilities of exclusion via simulations of LV dynamics and shape-based calculations"(Supplementary figures S3-S5), "Testing the dependence of the asymmetry index on size" (Supplementary figure S6), "Species information for the empirical example in Caracoles Ranch" (Supplementary tables S1-S2 and Supplementary figures S7-S11), "Estimation of the interaction matrices and intrinsic growth rates in Caracoles Ranch (Spain)", "Testing the diagonal-stability of Caracoles Ranch's interaction matrices", "Sampling coverage of Caracoles Ranch (Spain)"(Supplementary figure S12), "Results for Caracoles Ranch models" (Supplementary tables S3-S13 and Supplementary figures S13-S15), and "Results for bipartite plant-pollinator networks in Mangal" (Supplementary table S14 and Supplementary figures S17-S18).

### 1 Dependence of shape on size for feasibility domains

To illustrate the relation between shape and size when characterizing feasibility domains, here we review a counterexample that shows that the independence of both features is false.

Let us consider the simple case of communities with $S = 3$ species. As shown in Fig. 1, in such communities the spaces of growth rates can be represented by an sphere and the growth rates of stable coexistence (feasibility domains) by points that lie within spherical triangles. The border of those triangles limits the corresponding feasibility domains.

To highlight the relation between shape and size of feasibility domains with $S = 3$, let us now consider the area of a spherical triangle with angles $\alpha$, $\beta$, and $\gamma$ (measured in radians at the vertices, along the surface of the sphere). It is given by

$$\text{Area} = R^2 (\alpha + \beta + \gamma - \pi) = R^2 E, \tag{S1}$$

where $R$ is the radius of the sphere, and $E$ is called the spherical excess. In the case of spheres with $R = 1$ (like the ones considered in this paper), we can verify that tiny feasible domains (that is, feasible domains where Area $\to 0$) behave like planar triangles (where $E \to 0$) that meet the well-known relation: $\alpha + \beta + \gamma \to \pi$. However, if their area is large, then $\alpha + \beta + \gamma > \pi$. As an example, consider the case of a symmetrical feasible domain, where $\alpha = \beta = \gamma$ (see Fig. S1). According to Eq. (S1), if $R = 1$, then $\alpha = (\text{Area} + \pi)/3$. When Area $\to 0$ our feasible domain looks like a planar equilateral triangle ($\alpha = \pi/3$), whereas if its area is equal to one eighth of that of a full sphere (i.e., Area $= \pi/2$), then our equilateral triangle is an octant of a sphere ($\alpha = \pi/2$). That simple example shows that shape is not independent from size in spherical geometry.

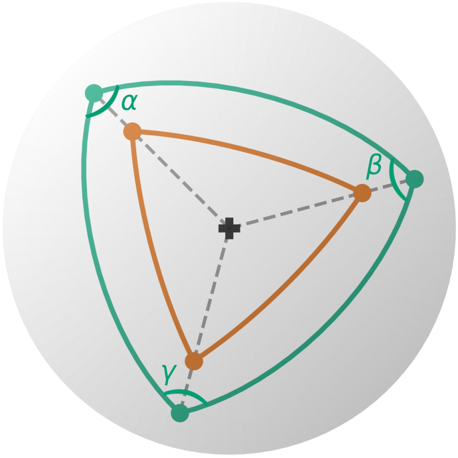


Fig. S1: An example of equilateral spherical triangles. Continuous lines show the border of the respective triangles, grey dashed lines represent the segments of great circles that connect their incenter (black plus marker) with the vertices of the triangles (orange and green dots, respectively). The area of the green triangle is 1.571 square units and their angles are equal to $\pi/2$ radians, whereas the area of the smaller triangle is 0.511 square units and their angles are $2\pi/5$ radians.

#### 2 Calculation of the incenter and the maximum isotropic perturbation for a given feasibility domain

Let us consider a community with $S$ species and an interaction matrix $\mathbf{A}$, that follows Eq. (1). To determine the maximum hyper-spherical cap $\kappa(\mathbf{A})$ that is tangent to all the sides of the feasibility domain's border and whose apex is the incenter $\mathfrak{I}$, we need to know the $S$ vertices and edges $\varepsilon^S_{S-1}$ (sections of great circles) that delimit the intersection of the feasibility domain with the unit-ball in $S$-dimensions, where $\varepsilon^S_{S-1}$ denotes the $S-1$-combination of a set $S$. Firstly, we define vertex $V_i$ as the point on the unit-ball where $N_i^* > 0$ and $N_j^* = 0$ for $j \neq i$. The structural vector $\mathbf{r}$ for vertex $V_i$ can be calculated as $\mathbf{v}_i = \mathbf{u}_i / \|\mathbf{u}_i\|$, where $\mathbf{u}_i = -\mathbf{A}\mathbf{e}_i$, $\mathbf{e}_i$ is a column vector whose $i$-th component is equal to 1 and the rest are 0, and $\|\mathbf{x}\| = \sqrt{x_1^2 + \cdots + x_S^2}$ represents the Euclidean norm of $\mathbf{x}$. Once we obtained the vertices of the feasibility domain, it is possible to derive the normal direction of the hyper-planes that contain the spherical borders of the feasibility domain, $\mathbf{n}_i$. To calculate the normal vectors $\mathbf{n}_i$, we use the generalization for the cross product in $S$ dimensions of $S-1$ vectors, which produces a vector that is perpendicular to those $S-1$ vectors. Here, we can express $\mathbf{n}_i$ as $\mathbf{n}_i = \mathbf{m}_i / \|\mathbf{m}_i\|$, where:

$$\mathbf{m}_i = \bigwedge_{\substack{k \neq i \\ k \in \{1, \cdots, S\}}} \mathbf{v}_k = \begin{vmatrix} v_1^1 & \cdots & v_1^S \\ \vdots & \ddots & \vdots \\ v_{S-1}^1 & \cdots & v_{S-1}^S \\ \mathbf{e}_1 & \cdots & \mathbf{e}_S \end{vmatrix}. \tag{S2}$$

Since normal vectors have 2 possible orientations, to place the incenter in the feasibility domain, for the normal vectors $\mathbf{n}_i$ we select the orientation that meets the condition $\mathbf{v}_i \cdot \mathbf{n}_j = \sum_{k=1}^{S} v_i^k n_j^k > 0$ for $i, jk \in \{1, \cdots, S\}$.

By using the normal vectors $\mathbf{n}_i$ calculated above, the incenter $\mathfrak{I}$ and its great circle distance to the boundary of the feasibility domain $\theta$ (denoted in the main text as the maximum isotropic perturbation) satisfy the following condition:

$$\begin{pmatrix} \mathbf{n}_1 \\ \hline \vdots \\ \hline \mathbf{n}_S \end{pmatrix} \mathfrak{I} = \begin{pmatrix} n_1^1 & \cdots & n_1^S \\ \vdots & \ddots & \vdots \\ n_S^1 & \cdots & n_S^S \end{pmatrix} \mathfrak{I} = \sin\theta \begin{pmatrix} 1 \\ \vdots \\ 1 \end{pmatrix}, \tag{S3}$$

which is given by the geometrical construction depicted in Fig. S2. Since $\mathfrak{I} \cdot \mathfrak{I} = 1$, and

$$\mathfrak{I} = \sin\theta \begin{pmatrix} n_1^1 & \cdots & n_1^S \\ \vdots & \ddots & \vdots \\ n_S^1 & \cdots & n_S^S \end{pmatrix}^{-1} \begin{pmatrix} 1 \\ \vdots \\ 1 \end{pmatrix}, \tag{S4}$$

the exact values of $\mathfrak{I}$ and $\theta$ can be obtained.

In addition, the position of a tangent point shared between the spherical cap (whose incenter is $\mathfrak{I}$) and the border of the FD located in the plane with normal vector $\mathbf{n}_i$, denoted as $\mathbf{T}_i$ (see Supplementary Fig. S2), can be calculated as $\mathbf{T}_i = \mathfrak{I} - \sin(\theta)\mathbf{n}_i$. The tangent point formula is derived by finding the point on the plane with normal vector $\mathbf{n}_i$ whose great circle distance to $\mathfrak{I}$ is $\theta$. That means that the Euclidean distance $d$ from $\mathfrak{I}$ to $\mathbf{T}_i$ is

$$d = \sqrt{\sum_{k=1}^{S} (I_k - T_{i,k})} = \sqrt{2 - 2\cos(\theta)}, \tag{S5}$$

where $I_k$ and $T_{i,k}$ represent the $k$-th component of $\mathfrak{I}$ and $\mathbf{T}_i$, respectively. Then, the components of $\mathbf{T}_i$ can be obtained by minimizing the function $f(\mathbf{T}_i) = d^2 - 2 + 2\cos(\theta)$ subject to the equality constraint $\mathbf{T}_i \cdot \mathbf{n}_i = 0$, with the method of the Lagrange multipliers.

Finally, note that it is possible to calculate the relative volume of the unit-ball covered by the spherical cap that is tangent to all the sides of the feasibility domain's border and whose apex is the incenter $\mathfrak{I}$ (see blue spherical caps in Fig. 1). That relative volume is given by:

$$\Xi(\mathbf{A}) = \frac{\text{area}\,(\kappa(\mathbf{A}))}{\text{area}\,(B^S)} = \frac{1}{2} I_{\sin^2\theta}\left(\frac{S-1}{2}, \frac{1}{2}\right), \tag{S6}$$

where $I_z\,(a, b)$ is the regularized incomplete beta function (Li, 2010). That relative volume can be used to compare the level of asymmetry between communities with the same number of species

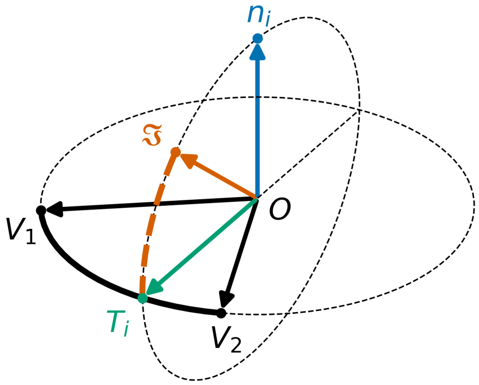


Fig. S2: 3D-representation of the relations between the incenter $\mathfrak{I}$ and a given side $\overline{V_1V_2}$ of the feasibility domain of a 3-species community. The black arrow represents the position of the vertices of the side$\overline{V_1V_2}$, the orange arrow the position of the incenter, the blue arrow the normal vector to the plane that contains the origin (O) and the vertices, and the green arrow the position of the point in the side $\overline{V_1V_2}$ that is tangent to $\mathfrak{I}$, $\mathbf{T}_i$. Finally, the orange dashed arc represents the great circle distance $\theta$ between the incerter $\mathfrak{I}$ and side $\overline{V_1V_2}$, that is, the arc $\mathfrak{I}\mathbf{T}_i$.

$(S_1 = S_2)$ and equal relative volumes $(\Omega_1 = \Omega_2)$, since it is closely related to the maximum isotropic perturbation $\theta$ that we introduced in the main text.

#### 3 Calculation of the exclusion distance of a given species

Following the notation and arguments introduced in the main text and Suppl. Section 2, the exclusion distance of species $i$, that is, the great circle distance between an initial vector of growth rates $\mathbf{r}_0$ and the border $\varepsilon_i$ of the feasibility domain where species $i$ goes excluded, is given by

$$\theta_{0,i} = \sin^{-1}\left(\mathbf{n}_i \cdot \mathbf{r}_0\right), \tag{S7}$$

where $\sin^{-1}(x)$ is the arcsine of $x$. Note that we simply replaced the incenter $\mathfrak{I}$ in Eqs. (S3) and (S4) by $\mathbf{r}_0$. Then, the position of the tangency point on the edge of the feasibility domain's border $\varepsilon_i$ (where species $i$ goes excluded) can be calculated as

$$\mathbf{T}_{0,i} = \mathbf{r}_0 - \sin(\theta_{0,i})\mathbf{n}_i. \tag{S8}$$

To conclude, we relate the point of tangency to the border of the feasibility domain with the minimum exclusion distance described in the main text. If we define the *isotropic feasible domain* of a vector $\mathbf{r}_0$ as the largest set of feasible vectors of intrinsic growth rates that can be achieved by perturbing isotropically (that is, in all directions) the vector $\mathbf{r}_0$, then such domain can be represented as the $S$-hyperspherical cap that meets the following conditions: i) its apex is $\mathbf{r}_0$, ii) it is tangent to the border of the feasibility domain (at point $\mathbf{r}_{0,\min}$), and iii) has no other points outside the feasibility domain, except those of tangency (see Fig. 2(a) for an example). As pointed out in our theoretical framework (see main text), species exclusion distance $\theta_0$ is the great circle distance between $\mathbf{r}_0$ and the point of tangency $\mathbf{r}_{0,\min}$). That point of tangency to the border of the feasibility domain, $\mathbf{r}_{0,\min}$, is just $\mathbf{r}_{0,\min} = \mathbf{T}_{0,q}$ with $\theta_{0,q} = \min(\theta_{0,i})$ for $i \in \{1, \cdots, S\}$, where $\theta_{0,q} = \theta_0$ defines the size of the minimum perturbation needed to exclude the one species.

### 4 Alternative definitions of the exclusion ratios

The definition of the exclusion ratio presented in the main text (Eq. (5)) can be generalized by replacing the term $1/S$ (i.e., the symmetric probability of exclusion) with a probability of exclusion derived from a null model approach:

$$ER_i(\mathbf{A}) = \frac{P_i^E(\mathbf{A})}{P_i^E(\mathbf{A}^{\text{Null}})}, \tag{S9}$$

where $\mathbf{A}^{\text{Null}}$ represents a random interaction matrix constructed by applying a null model to matrix $\mathbf{A}$.

### 5 Estimation of the probabilities of exclusion via simulations of LV dynamics and shape-based calculations

We can compute efficiently the probabilities of exclusion $P_i^E$ by applying the same quasi-Monte Carlo method used to estimate Eq. (3) to both the numerator and denominator in Eq. (6). The results rendered by that approach are in excellent agreement with those obtained from the simulation of the LV model (Eq. 1). The reason is that, given an initial, random vector of growth rates ($\mathbf{r}_0$) within the feasibility domain, the trend of species' abundances described by LV dynamics shows that the first species to be excluded matches that of our method, when the species' initial abundances are those of equilibrium ($\mathbf{N}^*$) and $\mathbf{r}_0$ is replaced by the vector of growth rates of the closest point in the border of the feasibility domain ($\mathbf{r}_{0,\min}$) (see panels (b) and (c) in Fig. 2 and Suppl. Fig. S3 for an example).

Nevertheless, during the preparation of this work, we have verified that the structure of the interactions (represented by the interaction matrix $\mathbf{A}$) influences the uncertainty of the above calculations. First, the smaller the size of the feasibility domain (or its relative volume $\Omega(\mathbf{A})$), the more difficult it is to randomly sample. Since the average probabilities of exclusion $P_i^E$ depend on that sampling, the smaller the feasibility domain, the greater the uncertainties of the corresponding sampling, and, hence, the larger those of the probabilities of exclusion of such domains. Second,

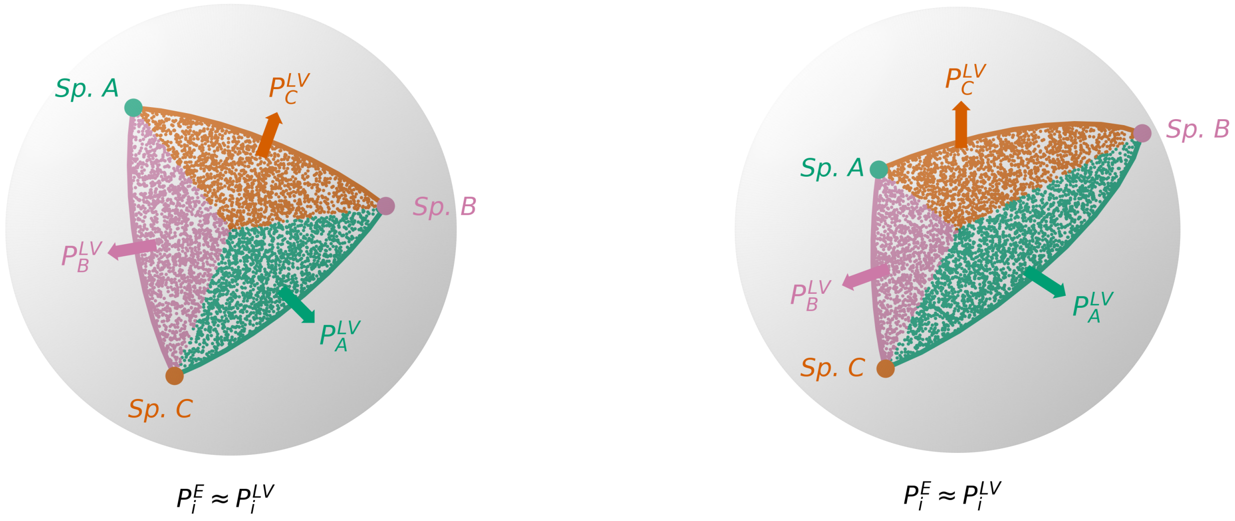


(a) (b)

Fig. S3: (a) Simulation of the first species to be excluded for 5,000 random initial, intrinsic growth rates ($\mathbf{r}_x$) within the feasibility domain shown in Fig. 2(b) by solving the LV equations (see Eq. (1)) when the species' initial abundances are those of equilibrium ($\mathbf{N}^*$) and the initial growth rates $\mathbf{r}_0$ are changed by those of the closest point of tangency in the border of the feasibility domain ($\mathbf{r}_{x,T}$). Colors of points are those of the first species that will be excluded. (e) Same results as in panel (a) for the feasibility domain in Fig. 2(b).

the more asymmetric the feasibility domain, the more difficult it is to sample the relative volumes of those species with a lower probability of exclusion (i.e., the more difficult it is to sample species that occupy smaller areas in figures analogous to our Fig. 2(c)). Thus, the exclusion probabilities of the potentially more persistent species (those that will have a smaller relative volume within the feasibility domain) will be more uncertain than the exclusion probabilities of the less persistent species (since the latter are easier to randomly sample). Finally, we observed that the differences between LV estimations and those provided by shape-based calculations increase with asymmetry (see Suppl. Fig. S4). However, our tests with symmetric feasibility domains suggest that shape-based calculations are more accurate (see the example for asymmetry index $J'(\mathbf{A}) = 1$ in Suppl. Fig. S4).

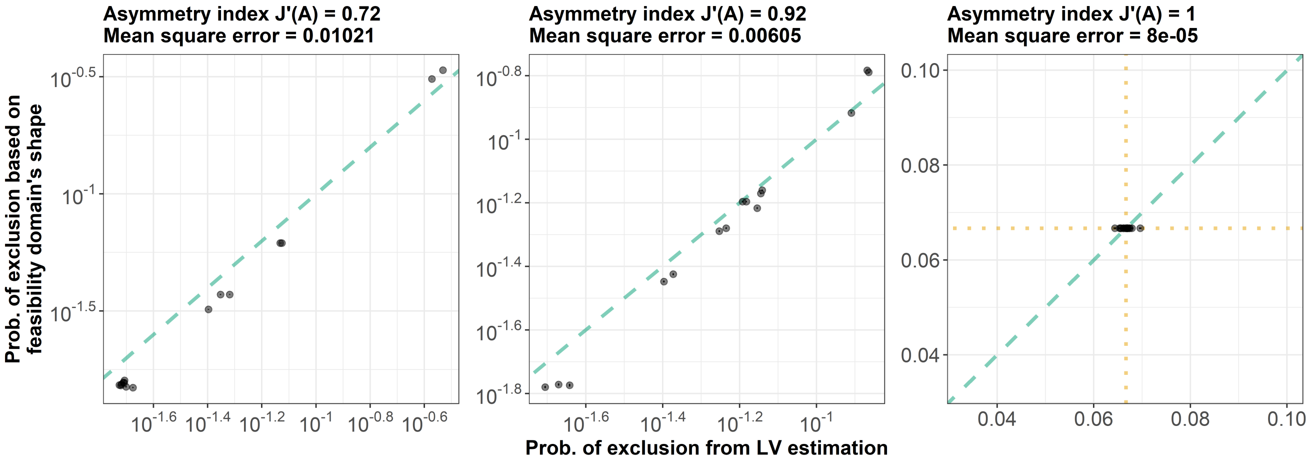


Fig. S4: Comparison between the probabilities of exclusion $P_i^E(\mathbf{A})$ obtained from LV simulations and those estimated with Eqs. (3) and (6). Each panel represents the results a community with $S = 15$ species. The value of the asymmetry index and of the mean square error between the set of probabilities of exclusion are shown in each panel's title. Green dashed line is a guide to the eye that locates probabilities of exclusion that are equal under both calculation approaches, whereas orange dotted line shows the theoretical value of $P_i^E(\mathbf{A})$ for symmetrical feasibility domains, i.e., $P_i^E(\mathbf{A}) = 1/S$ for $J'(\mathbf{A}) = 1$.

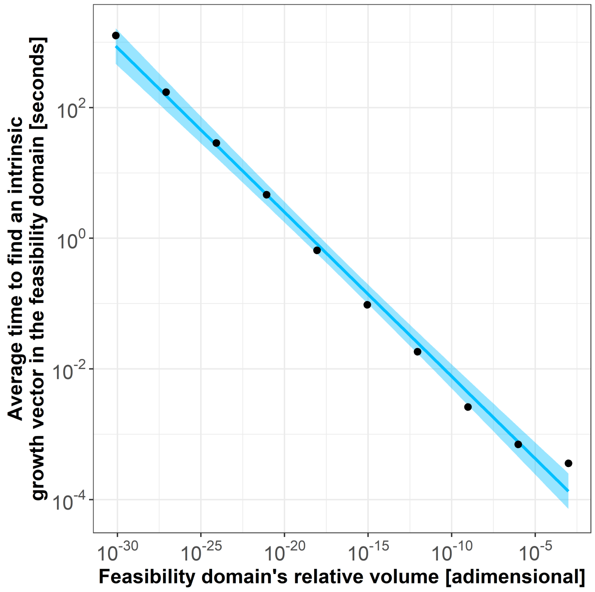


Fig. S5: Dependence of the average computing time (in seconds) for randomly finding feasible intrinsic growth rates in symmetric communities on the feasibility domain's relative volume, $\Omega(\mathbf{A})$ (see Eq. (2)). We used the algorithm "random_feasible_growth_rates" available at `https://github.com/RadicalCommEcol/Asymmetric_Interactions/R/6_computing_times_LV.R`. To test our algorithm we used interaction matrices $\mathbf{A} = -I_S$, where $I_S$ represents a $S \times S$ identity matrix, and $S$ takes the following values: 10, 20, 30, 40, 50, 60, 70, 80, 90 and 100. Blue line shows the fitted regression model computing time $= 10^{-4.62889}\Omega^{-0.25147}$, whereas the blue shade area shows the 95% confidence interval of such model.

### 6 Testing the dependence of the asymmetry index on size

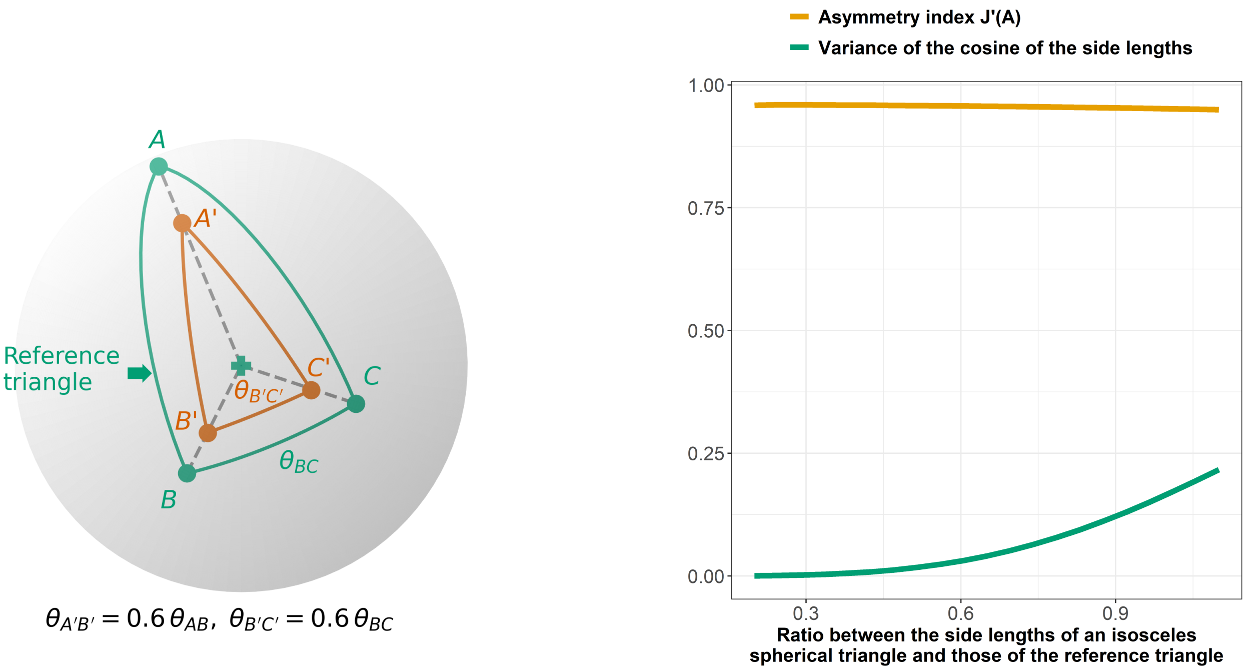


(a) (b)

Fig. S6: (a) An example of (isosceles) spherical triangles with proportional sides. Continuous lines show the border of the respective triangles, dashed lines show the segments of great circles that connect the incenter of the reference triangle (black plus marker) with the vertices of the triangles (orange and green dots, respectively). The ratio between the sides of the orange triangle and those of the green one is equal to 0.6. (b) Comparison of shape indices for isosceles spherical triangles whose side lengths are proportional, like those in Fig. S6(a). The horizontal axis shows the ratio between the side lengths of the spherical triangles tested and those of the reference triangle. Results for the shape index proposed in (Grilli *et al.*, 2017), the variance of the cosine of side lengths, are in green; whereas those for $J'(\mathbf{A})$, obtained with Eq. (7), are in orange.

### 7 Species information for the empirical example in Caracoles Ranch (Spain)

Table S1: Species studied and their overall abundance (individuals counted) in the whole study area for each year of observations.

| Species | Family | 2015 | 2016 | 2017 | 2018 | 2019 | 2020 | 2021 |
|---|---|---|---|---|---|---|---|---|
| *Beta macrocarpa* | Amaranthaceae | 136 | 1,230 | 90 | 20 | 1,052 | 2,023 | 459 |
| *Centaurium tenuiflorum* | Gentianaceae | 1,206 | 37 | 1,057 | 522 | 705 | 1,946 | 1,603 |
| *Chamaemelum fuscatum* | Asteraceae | 5,050 | 5,802 | 2,619 | 0 | 3,989 | 1,348 | 1,451 |
| *Chamaemelum mixtum* | Asteraceae | 882 | 909 | 208 | 2 | 9 | 144 | 68 |
| *Hordeum marinum* | Poaceae | 6,719 | 13,159 | 54,264 | 43,355 | 71,923 | 16,780 | 16,465 |
| *Lythrum tribracteatum* | Lythraceae | 1,321 | 474 | 676 | 3 | 0 | 0 | 0 |
| *Leontodon maroccanus* | Asteraceae | 2,485 | 3,448 | 21,657 | 814 | 4,182 | 9,098 | 11,455 |
| *Melilotus elegans* | Fabaceae | 313 | 135 | 2 | 0 | 6 | 1 | 31 |
| *Melilotus sulcatus* | Fabaceae | 504 | 414 | 36 | 1 | 1,059 | 1,106 | 2,418 |
| *Parapholis incurva* | Poaceae | 1,239 | 78 | 2,980 | 29 | 7,534 | 886 | 678 |
| *Plantago coronopus* | Plantaginaceae | 5,439 | 518 | 2,427 | 1 | 118 | 1,134 | 321 |
| *Polypogon maritimus* | Poaceae | 4,101 | 709 | 5,085 | 315 | 7,332 | 3,367 | 25 |
| *Polypogon monspeliensis* | Poaceae | 442 | 26 | 89 | 1 | 284 | 427 | 78 |
| *Pulicaria paludosa* | Asteraceae | 124 | 201 | 2,899 | 746 | 739 | 1418 | 0 |
| *Salsola soda* | Amaranthaceae | 7,346 | 490 | 1,629 | 4,771 | 791 | 2,222 | 2,821 |
| *Scorzonera laciniata* | Asteraceae | 881 | 226 | 85 | 17 | 793 | 1,334 | |
| *Sonchus asper* | Asteraceae | 237 | 41 | 0 | 1 | 366 | 1,021 | 842 |
| *Spergularia rubra* | Caryophyllaceae | 146 | 103 | 108 | 0 | 550 | 2,318 | 102 |
| *Suaeda splendens* | Amaranthaceae | 60 | 26 | 0 | 1 | 52 | 99 | 0 |

Table S2: Species studied and their average seed survival and germination rates.

| Species | Seed survival rate | Germination rate |
|---|---|---|
| *Beta macrocarpa* | 0.43 | 0.38 |
| *Centaurium tenuiflorum* | 0.10 | 0.53 |
| *Chamaemelum fuscatum* | 0.38 | 0.08 |
| *Chamaemelum mixtum* | 0.32 | 0.76 |
| *Hordeum marinum* | 0.25 | 0.94 |
| *Lythrum tribracteatum* | 0.51 | 0.62 |
| *Leontodon maroccanus* | 0.33 | 0.89 |
| *Melilotus elegans* | 0.60 | 0.59 |
| *Melilotus sulcatus* | 0.63 | 0.77 |
| *Parapholis incurva* | 0.60 | 0.33 |
| *Plantago coronopus* | 0.57 | 0.84 |
| *Polypogon maritimus* | 0.46 | 0.85 |
| *Polypogon monspeliensis* | 0.49 | 0.91 |
| *Pulicaria paludosa* | 0.55 | 0.84 |
| *Salsola soda* | 0.30 | 0.52 |
| *Scorzonera laciniata* | 0.41 | 0.69 |
| *Sonchus asper* | 0.29 | 0.60 |
| *Spergularia rubra* | 0.41 | 0.44 |
| *Suaeda splendens* | 0.58 | 0.61 |

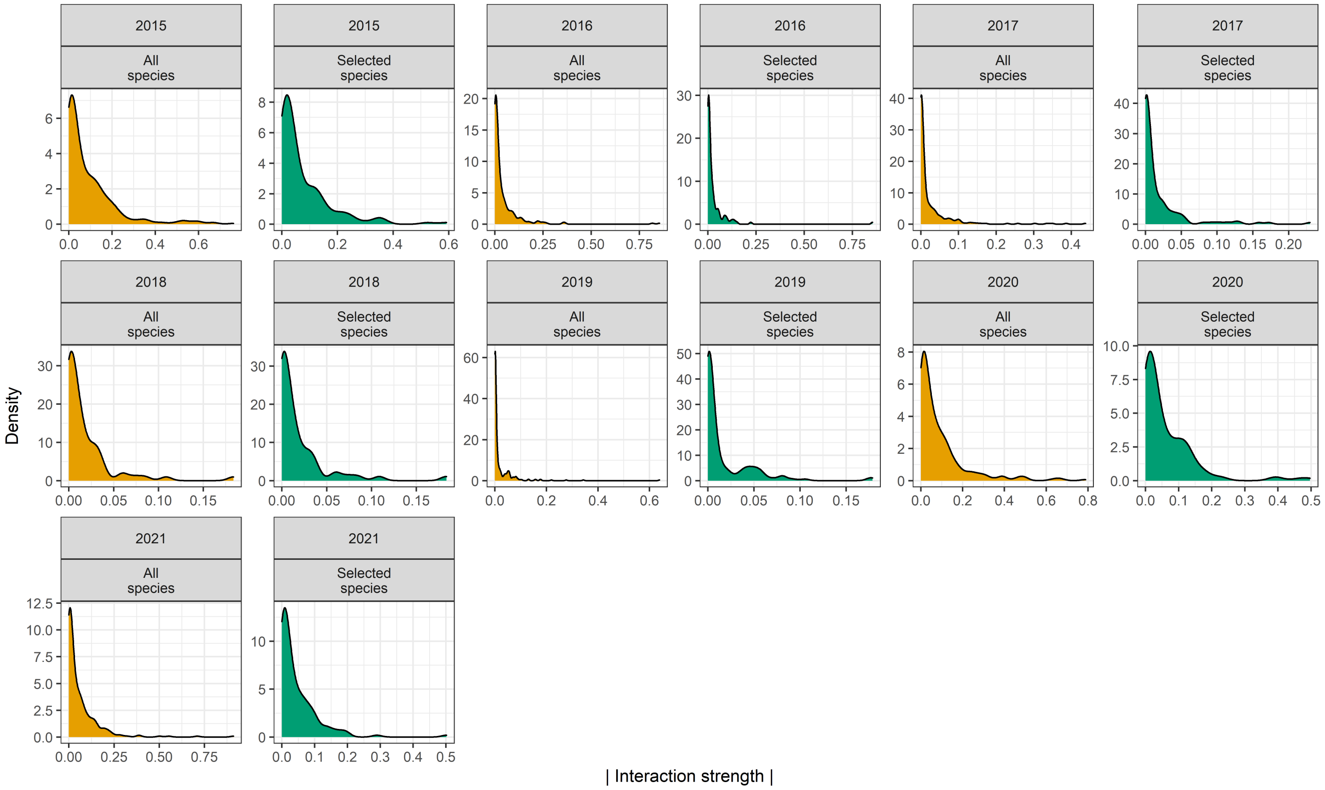


Fig. S7: Distribution of the absolute value of the interaction strengths in Caracoles Ranch across the sampling years. We present the results for the whole set of plant species that were present in a given year (orange fill) and for those seven species that were present all the years (green fill).

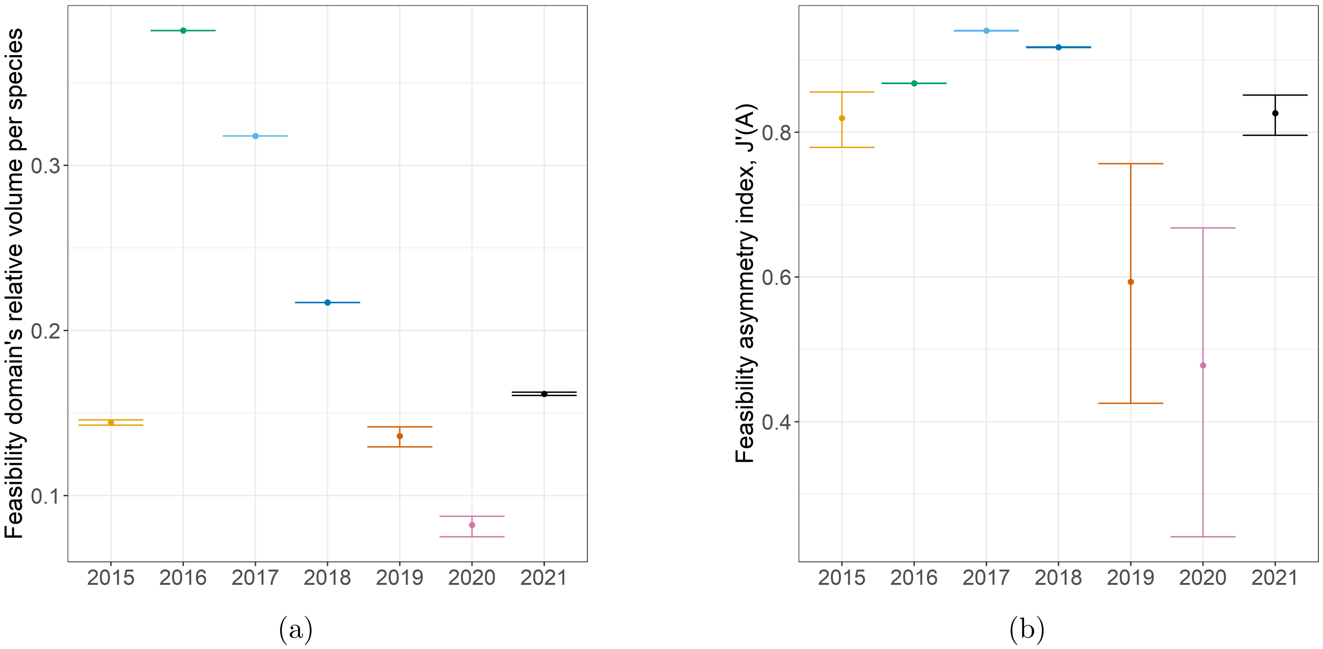


Fig. S8: (a) Time evolution of the feasibility domain's relative volume per species ($\Omega^{1/S} \in [0, 0.5]$, where $S$ is the total number of species (Song *et al.*, 2018) [see Eqs. (2) and (3)]) for Caracoles Ranch. (b) Asymmetry index ($J' \in [0, 1]$ [see Eq. (7)]) for the data in panel (a). Error bars in both panels show the 95% confidence intervals for the results of the 10,000 random draws generated to estimate the species' probability of exclusion ($P_i^E(\mathbf{A})$) each year, by using the quasi-Monte Carlo estimation related to Eq. (3).

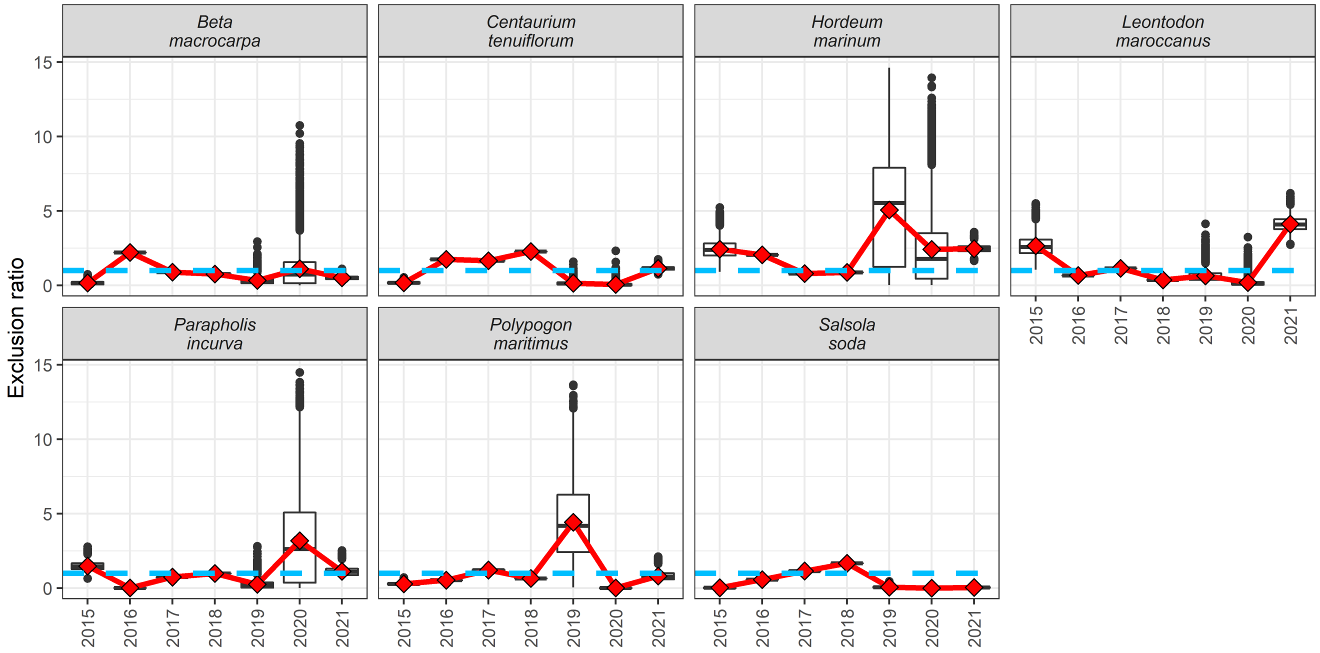


Fig. S9: Time evolution of the exclusion ratio $ER_i(\mathbf{A})$ for each plant species (see Eq. (5)). The box-plots show the results of the 10,000 random draws generated to estimate $P_i^E(\mathbf{A})$ generated each year, by using the quasi-Monte Carlo estimation related to Eq. (3). The red continuous lines connect the mean values of $ER_i(\mathbf{A})$ (red diamonds) from consecutive years, whereas the blue dashed line indicates the region where $ER_i = 1$ (i.e., $P_i^E(\mathbf{A}) = 1/S$). We only show the results for the seven plant species selected for the analyses (see main text).

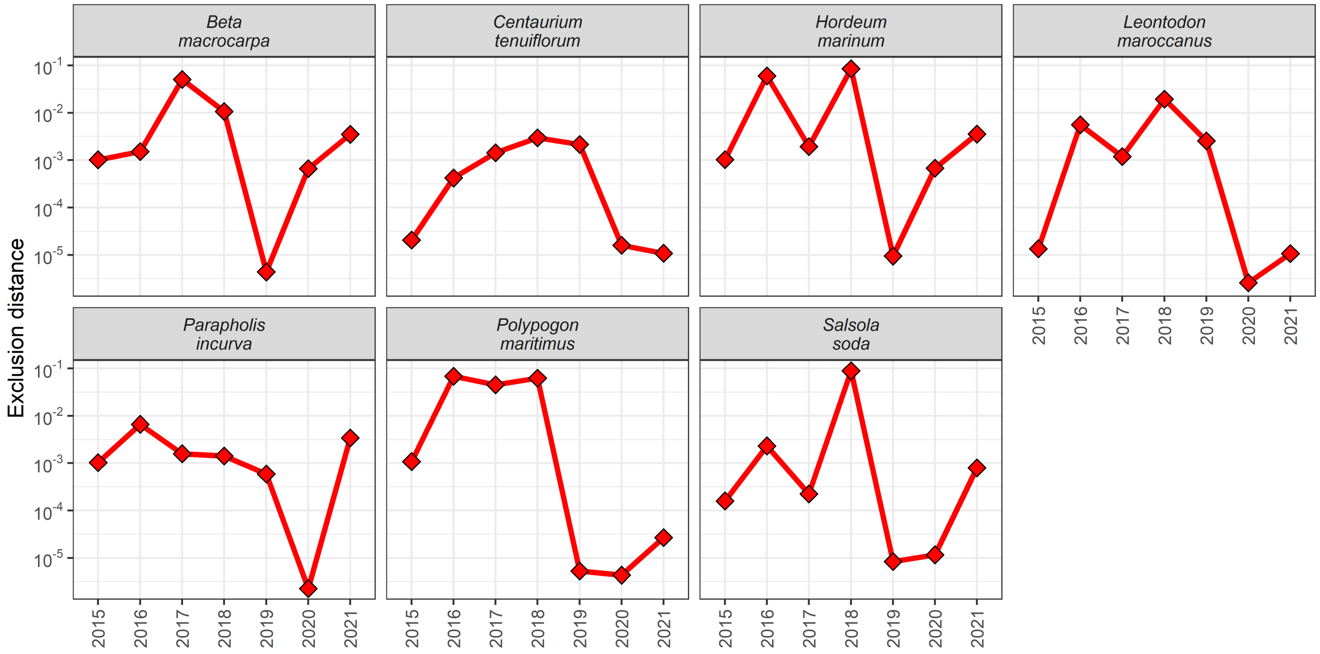


Fig. S10: Time evolution of exclusion distance. The red continuous lines connect the species values (red diamonds) from consecutive years. We only show the results for the seven plant species selected for the analyses (see main text).

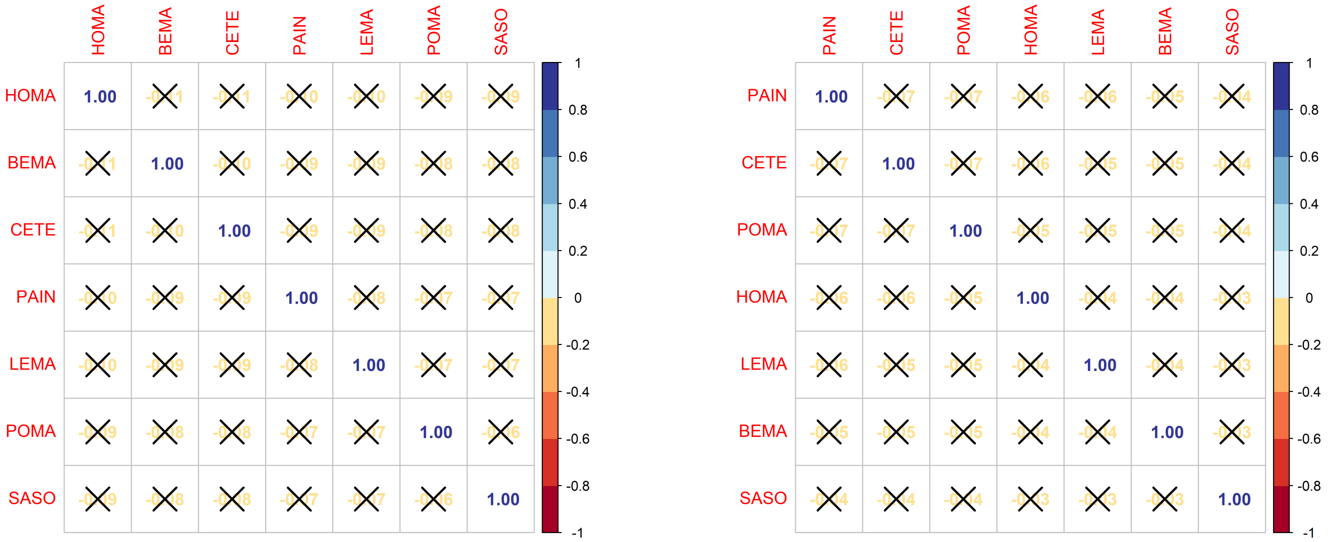


(a) (b)

Fig. S11: a) Correlogram for the time evolution of the mean value of the exclusion ratios shown in Fig. S9. b) Correlogram for the time evolution of the exclusion distances in Fig. S10. Crosses show correlations whose p-value > 0.05.

### 8 Estimation of the interaction matrices and intrinsic growth rates in Caracoles Ranch (Spain)

To estimate the pairwise per capita effects of plant species and the intrinsic growth rates for the grassland of Caracoles Ranch in a given year, we applied the methodology presented by García-Callejas *et al.* (2021). The parameterization is based on the Ricker model, and its equivalence to a linear LV model.

The elements of our interaction matrices $\mathbf{A}$ (that is, the per capita effect of species j over species i, $\alpha_{ij}$) were estimated following the methods in the section "Estimating species interaction networks and intrinsic growth rates" in García-Callejas *et al.* (2021). Indeed, the values we used here were those provided by the authors. The data were extracted from Zenodo (`https://doi.org/10.5281/zenodo.5390313`) and deposited in our repository at:

`Data/caracoles_raw_data/alpha_heterogeneous_time.csv`

Regarding the estimation of the vectors of intrinsic growth rates, we calculated an equivalent intrinsic growth rate, by following the Suppl. Note 2 in (García-Callejas *et al.*, 2021). Such value is given by:

$$r_i = \lambda_i + \ln\left(\frac{1-(1-g_i)s_i}{g_i}\right). \tag{S10}$$

for species $i$, where $s_i$ is the survival rate of seeds in the seed bank, $g_i$ is seed germination rate (see )), and $\lambda_i$ is the number of seeds produced by species $i$ in the absence of interactions. We used the empirical information in table S2 and the file `Data/caracoles_raw_data/lambda_heterogeneous_both.csv` (see our repository) to extract the value of the vector of growth rates $\mathbf{r}$ for a given year. Since our framework is developed for feasible equilibrium points, we checked the condition $\mathbf{N}^* = \mathbf{A}^{-1}\mathbf{r} > 0$. If we found a species with negative equilibrium abundance $N_i^* < 0$, we replaced that value by $N_i^{*,mod} = 1$; otherwise, we did nothing (that is, we set $N_i^{*,mod} = N_i^*$). Finally, recalculated the value of $\mathbf{r} = \mathbf{A}\mathbf{N}^{*,mod}$. The code we used can be accessed in our repository at: `R/1c_caracoles_calculate_arc_distances.R`

### 9 Testing the d-stability of Caracoles Ranch's interaction matrices

Testing whether the interaction matrices of our system are diagonally-stable (or d-stable) is equivalent to searching for a positive diagonal matrix $\mathbf{D}$ that meets the following condition: the eigenvalues of $\mathbf{W} = (\mathbf{A}^T)\mathbf{D} + \mathbf{DA}$ are negative, where $\mathbf{A}^T$ is the transpose of the interaction matrix $\mathbf{A}$ (Takeuchi, 1996). For each sampled year, we used the genetic algorithms in the R-library `GA` v3.2.1 (Scrucca, 2013, 2017), as well as the solver of non-linear equations in the R-library `nleqslv`. v3.3.2 Hasselman (2018), to minimize the maximum eigenvalue of $\mathbf{W}$ None of the matrices we found met the above condition. The code we used can be accessed in our repository at:

- `R/10_computing_diagonal_stability_Caracoles_GA.R`
- `R/10_computing_diagonal_stability_Caracoles_V2.R`

These complementary analyses showed that the interaction matrices of our annual grassland were not D-stable. Thus, we cannot ensure that dynamic stability is fulfilled in addition to feasibility in that system.

### 10 Sampling coverage of Caracoles Ranch (Spain)

We have estimated the sampling coverage of (individual) plant–plant interactions in each year, by pooling the data for all plant species, and using the coverage framework introduced in Chao *et al.* (2014). The sampling coverage (or coverage) of the observed interactions is simply their total relative abundances, or equivalently, the proportion of the total number of interactions in an assemblage that belong to interactions represented in the sample. To calculate the coverage in each plot, we used the R-package `iNEXT` v3.0.0 (Hsieh *et al.*, 2016). As can be seen in Fig. S12, the sampling coverage from 2015 to 2021 is approximately equal to 99%. This means that less than 1% of the total interactions in the assemblage belong to undetected interactions in a given year. For this reason, extremely rare, undetected interactions do not make a significant contribution to that proportion, even if there are many such interactions. In addition, note that the above estimations do not account for the existence of forbidden interactions, that is, non-occurrences of pairwise interactions that can be accounted for by biological constraints, such as spatio-temporal uncoupling, size or reward mismatching, foraging constraints and physiological–biochemical constraints (Jordano, 2016). Hence, the real sampling coverage of our plant-plant interactions could be higher than 99%.

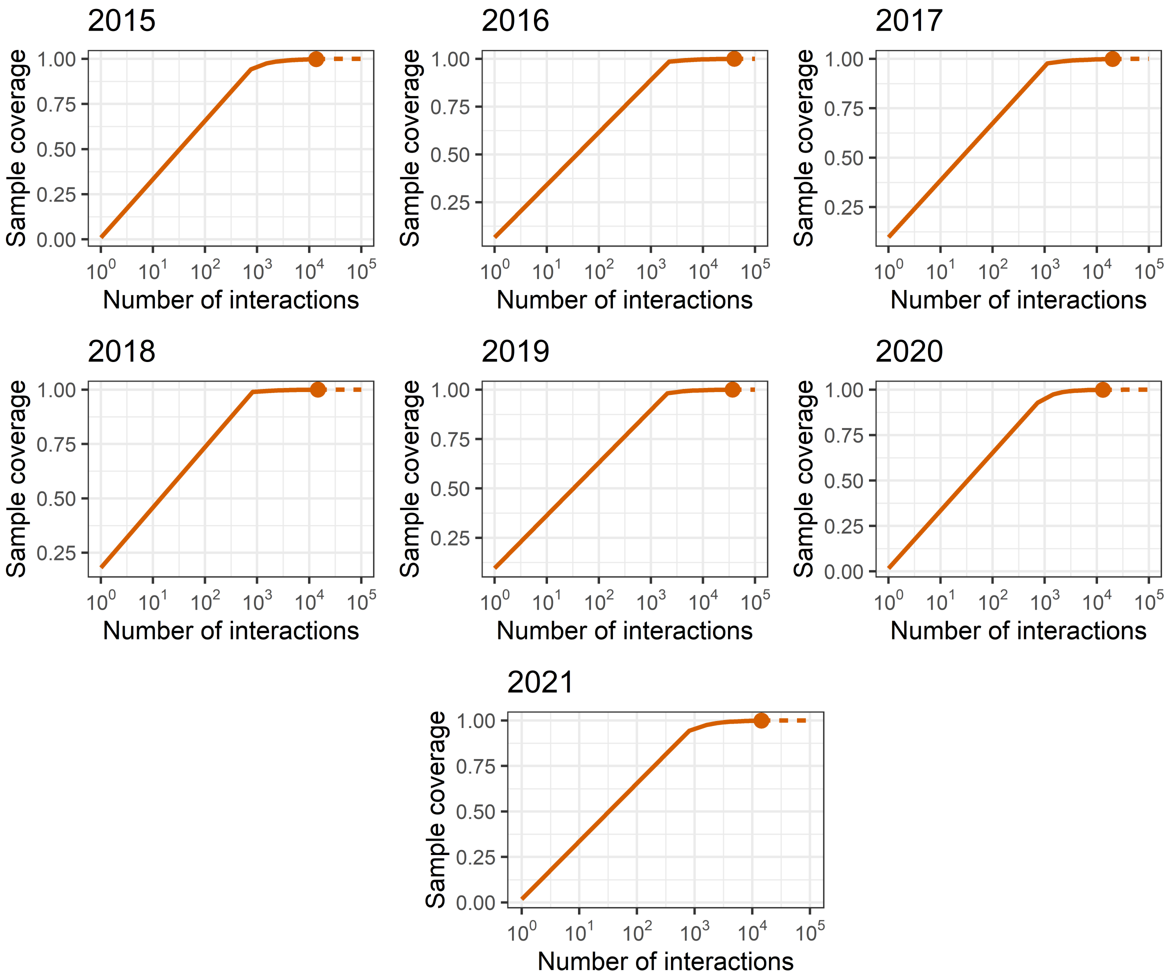


Fig. S12: Sample coverage for rarefied samples (solid line) and extrapolated samples (dashed line) as a function of sample size for plant-plant interactions sampled from each annual community. The 95% confidence intervals were obtained by a bootstrap method based on 1,000 replications. Actual sampling coverage is denoted by solid dots.

### 11 Results for Caracoles Ranch models

Table S3: Results for the Gamma-GLM (with log link function) showing the effect of the exclusion ratio in a given year $t$ on the population growth $N_{i,t+1}/N_{i,t}$. We highlighted in bold the most important estimates based on effect sizes and its variability. P-values have been obtained from the summary report on the models fitted with R-library `stats` v4.1.0 (R Core Team, 2021).

| Population growth | Estimate | Std. Error | t value | Pr($>\|z\|$) |
|---|---|---|---|---|
| **(Intercept)** | **3.0922** | **0.8101** | **3.817** | **0.000563** |
| Excl. ratio | 1.0710 | 0.9669 | -1.108 | 0.275996 |

Table S4: Results for the Gamma-GLM (with log link function) showing the effect of the probability of exclusion in a given year $t$ on the population growth $N_{i,t+1}/N_{i,t}$. We highlighted in bold the most important estimates based on effect sizes and its variability. P-values have been obtained from the summary report on the models fitted with R-library `stats` v4.1.0 (R Core Team, 2021).

| Population growth | Estimate | Std. Error | t value | Pr($>\|z\|$) |
|---|---|---|---|---|
| **(Intercept)** | **1.968** | **0.690** | **2.853** | **0.00743** |
| Prob. of excl. | 6.625 | 5.765 | 1.149 | 0.25874 |

Table S5: Results for the Gamma-GLM (with log link function) showing the effect of the exclusion distance in a given year $t$ on the population growth $N_{i,t+1}/N_{i,t}$. We highlighted in bold the most important estimates based on effect sizes and its variability. P-values have been obtained from the summary report on the models fitted with R-library `stats` v4.1.0 (R Core Team, 2021).

| Population growth | Estimate | Std. Error | t value | Pr($>\|z\|$) |
|---|---|---|---|---|
| **(Intercept)** | **2.685** | **0.592** | **4.535** | **7.22e-05** |
| Excl. distance | -10.860 | 20.946 | -0.518 | 0.608 |

Table S6: Results for the Gaussian-GLM (with log link function) showing the effect of abundance in a given year $t$ on the population growth $N_{i,t+1}/N_{i,t}$. We highlighted in bold the most important estimates based on effect sizes and its variability. P-values have been obtained from the summary report on the models fitted with R-library `stats` v4.1.0 (R Core Team, 2021).

| Population growth | Estimate | Std. Error | t value | Pr($>\|z\|$) |
|---|---|---|---|---|
| **(Intercept)** | **5.181539** | **0.336858** | **15.382** | **¡2e-16** |
| Abundance | -0.016842 | 0.009603 | -1.754 | 0.0887 |

Table S7: Results for the Gaussian-GLM (with log link function) showing the effect of the probability of exclusion in a given year $t$ on the population growth $N_{i,t+1}/N_{i,t}$. We highlighted in bold the most important estimates based on effect sizes and its variability. P-values have been obtained from the summary report on the models fitted with R-library `stats` v4.1.0 (R Core Team, 2021).

| Population growth | Estimate | Std. Error | t value | Pr(> $\|z\|$) |
|---|---|---|---|---|
| **(Intercept)** | **2.4874** | **0.8518** | **2.920** | **0.00627** |
| Prob. of excl. | 1.0508 | 6.2424 | 0.168 | 0.86734 |

Table S8: Results for the Gaussian-GLM (with log link function) showing the effect of the exclusion distance in a given year $t$ on the population growth $N_{i,t+1}/N_{i,t}$. We highlighted in bold the most important estimates based on effect sizes and its variability. P-values have been obtained from the summary report on the models fitted with R-library `stats` v4.1.0 (R Core Team, 2021).

| Population growth | Estimate | Std. Error | t value | Pr(> $\|z\|$) |
|---|---|---|---|---|
| **(Intercept)** | **2.6408** | **0.6108** | **4.324** | **0.000133** |
| Excl. distance | -5.9275 | 31.7624 | -0.187 | 0.853102 |

Table S9: Results for the non-linear least squares (NLS) fit of the model $\log_{10}(\text{Pop. growth} + 1) = a \cdot \exp(b \cdot \text{Excl. ratio})$, showing the effect of the exclusion ratio in a given year $t$ on the population growth $N_{i,t+1}/N_{i,t}$. We highlighted in bold the most important estimates based on effect sizes and its variability. P-values have been obtained from the summary report on the models fitted with R-library `stats` v4.1.0 (R Core Team, 2021).

| Population growth | Estimate | Std. Error | t value | Pr(> $\|z\|$) |
|---|---|---|---|---|
| **a** | **0.646** | **0.136** | **4.766** | **3.67e-05** |
| b | -0.318 | 0.3696 | -0.861 | 0.396 |

Table S10: Results for the GLMM model showing the effect of the exclusion ratio in a given year $t$ on $\log_{10}(\text{Pop. growth} + 1)$, where $N_{i,t+1}/N_{i,t}$ represents the population growth. The model uses plant species as a random factor and includes an auto-correlation term to control for temporal correlation. We highlighted in bold the most important estimates based on effect sizes and its variability. P-values have been obtained from the summary report on the model fitted with R-library `lme4` v1.1-28 (Bates *et al.*, 2015).

| Population growth | Estimate | Std. Error | t value | Pr(> $\|z\|$) |
|---|---|---|---|---|
| **(Intercept)** | **-1.2622388** | **0.3393160** | **-3.719951** | **0.0009** |
| Excl. ratio | 0.1318161 | 0.4050064 | -0.325467 | 0.7473 |

Table S11: Results for the non-linear least squares (NLS) fit of the model $\log_{10}(\text{Pop. growth} + 1) = a \cdot \exp(b \cdot \text{Prob. of exclusion})$, showing the effect of the probability of exclusion in a given year $t$ on the population growth $N_{i,t+1}/N_{i,t}$. We highlighted in bold the most important estimates based on effect sizes and its variability. P-values have been obtained from the summary report on the models fitted with R-library `stats` v4.1.0 (R Core Team, 2021).

| Population growth | Estimate | Std. Error | t value | Pr(> $\|z\|$) |
|---|---|---|---|---|
| **a** | **0.5926** | **0.1422** | **4.168** | **0.0002** |
| b | -0.800 | 2.222 | -0.360 | 0.721 |

Table S12: Results for the non-linear least squares (NLS) fit of the model $\log_{10}(\text{Pop. growth} + 1) = a \cdot \exp(b \cdot \text{Excl. distance})$, showing the effect of the exclusion distance in a given year on the population growth $N_{i,t+1}/N_{i,t}$. We highlighted in bold the most important estimates based on effect sizes and its variability. P-values have been obtained from the summary report on the models fitted with R-library `stats` v4.1.0 (R Core Team, 2021).

| Population growth | Estimate | Std. Error | t value | Pr(> $\|z\|$) |
|---|---|---|---|---|
| **a** | **0.5312** | **0.1074** | **4.945** | **2.17e-05** |
| b | 3.3297 | 5.8291 | 0.571 | 0.572 |

Table S13: AIC comparison among the candidate models for the population growth $N_{i,t+1}/N_{i,t}$ in Caracoles Ranch (see main text). In this table, $\text{AIC}_i$ is the AIC of the $i$-th model, $\text{AIC}_{\text{min}}$ is the lowest AIC one obtains among the set of models examined, and $\Delta_i$ is given by $\Delta_i = \text{AIC}_i - \text{AIC}_{\text{min}}$. The following models have not been considered in this table because they did not converge: NLS (exp.) model for abundance, GLM (Gamma with log link) for abundance, and GLM (Gaussian with log link) for exclusion ratio.

| Model | Degrees of freedom | $\text{AIC}_i$ | $\Delta_i$ |
|---|---|---|---|
| NLS (exp.) for Excl. ratio | 3 | 63.86213 | 0 |
| NLS (exp.) for Excl. distance | 3 | 64.86826 | 1.00613 |
| NLS (exp.) for Prob. of exclusion | 3 | 65.02887 | 1.16674 |
| GLMM (exp.) for Excl. ratio | 5 | 139.22319 | 75.36106 |
| GLM (Gamma with log link) for Prob. of exclusion | 3 | 197.69274 | 133.8306 |
| GLM (Gamma with log link) for Excl. ratio | 3 | 197.71752 | 133.8554 |
| GLM (Gamma with log link) for Excl. distance | 3 | 198.27493 | 134.4128 |
| GLM (Gaussian with log link) for Abundance | 3 | 345.6220 | 281.7599 |
| GLM (Gaussian with log link) for Excl. distance | 3 | 369.9087 | 306.0466 |
| GLM (Gaussian with log link) for Prob. of exclusion | 3 | 369.9261 | 306.064 |

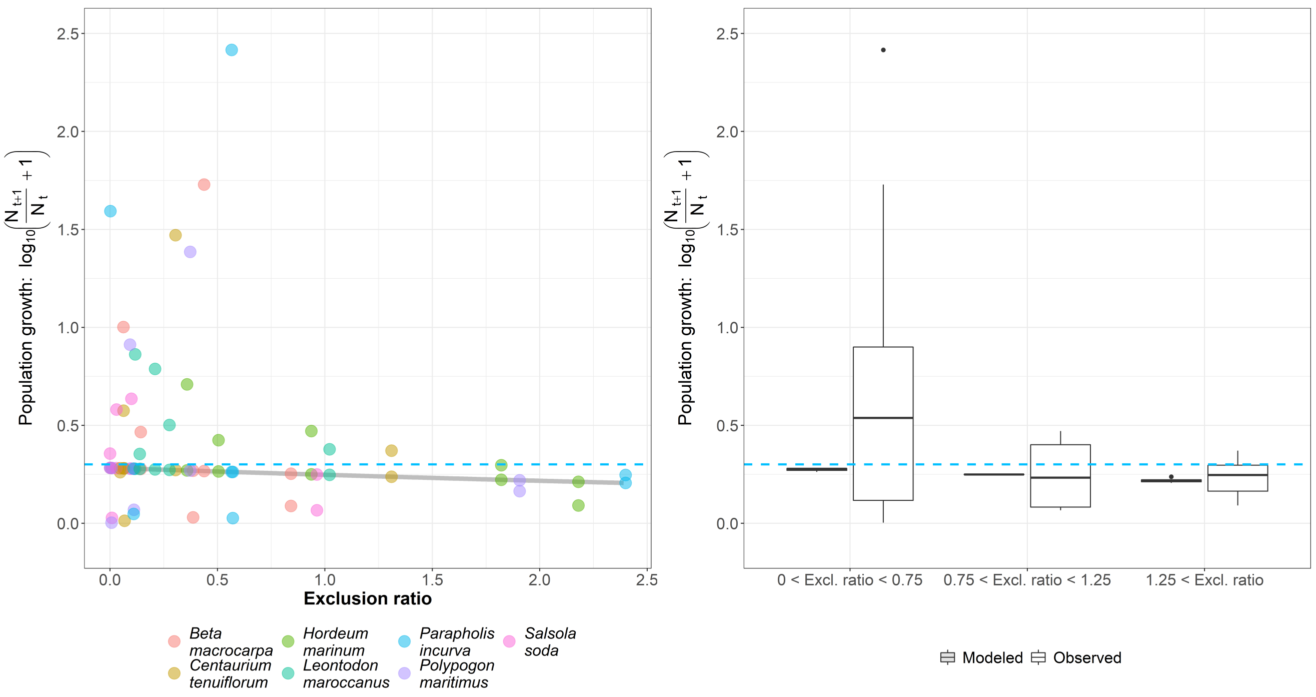


Fig. S13: Results for the GLMM in table S10 showing the effect of the exclusion ratio in a given year on observed population growth rate. Left panel shows the observed values of exclusion ratio and population growth for each studied plant species (colored dots), and the average value predicted by the GLMM model. Box-plots in the right panel display the observed and predicted population growth across three different groups of exclusion ratios: i) $ER_i < 0.75$, ii) $.75 \leq ER_i < 1.25$ and iii) $1.25 \leq ER_i$.

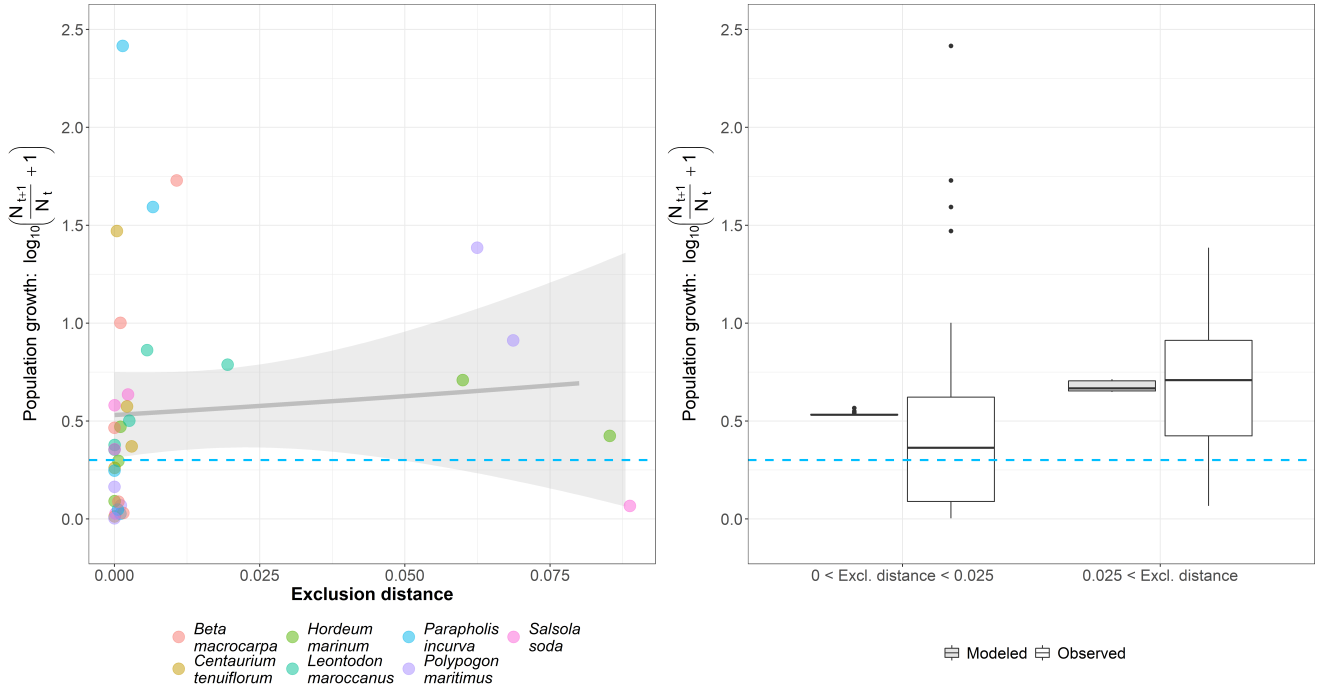


Fig. S14: Results for the non-linear model $\log_{10}(\text{Pop. growth} + 1) = a \cdot \exp(b \cdot \text{Excl. distance})$ in table S12, showing the effect of the exclusion distance in a given year $t$ on the population growth.Left panel shows the observed values of exclusion ratio and population growth for each studied plant species (colored dots), and the values predicted by the NLS model (grey line) and their 95% confidence interval (grey shaded area). Box-plots in the right panel display the observed and predicted population growth across three different groups of exclusion distances: i) $\theta_i < 0.025$, and ii) $0.025 \leq \theta_i$.

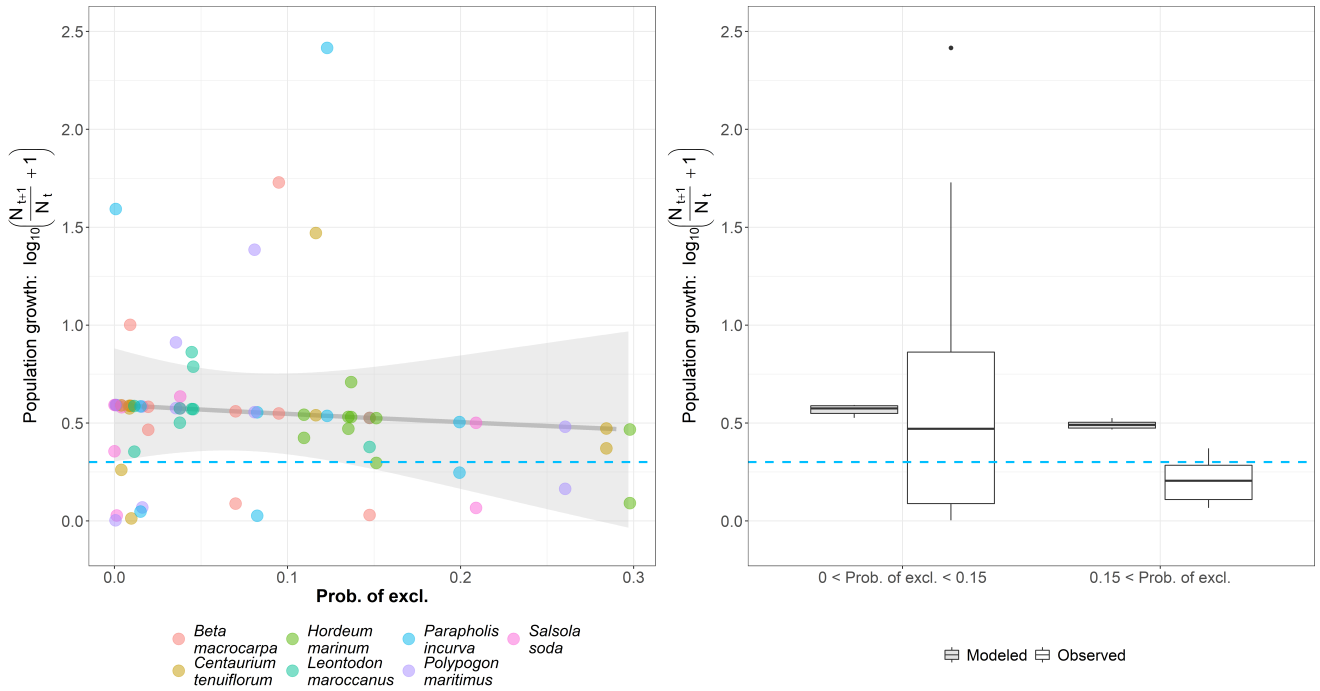


Fig. S15: Results for the non-linear model $\log_{10}(\text{Pop. growth} + 1) = a \cdot \exp(b \cdot \text{Prob. of excl.})$ in table S11, showing the effect of the probability of exclusion in a given year $t$ on observed population growth rate. Left panel shows the observed values of probability of exclusion and population growth for each studied plant species (colored dots), and the values predicted by the NLS model (grey line) and their 95% confidence interval (grey shaded area). Box-plots in the right panel display the observed and predicted population growth across two different groups of probability of exclusion: i) $P_i^E < 0.155$, and ii) $0.15 \leq P_i^E$.

### 12 Results for bipartite plant-pollinator networks in Mangal

Table S14: List of studies in Mangal database (Poisot *et al.*, 2016) ordered by author with the year of publication, number of contributed networks and digital object identifier (DOI).

| First author | Number of networks | Year | DOI |
|---|---|---|---|
| Bartomeus | 12 | 2008 | 10.1007/s00442-007-0946-1 |
| Bezerra | 1 | 2009 | 10.1111/j.1365-2656.2009.01567.x |
| CaraDonna | 25 | 2020 | 10.1111/ele.12740 |
| Kaiser-Bunbury | 1 | 2014 | 10.1890/14-0024.1 |
| Ponisio | 49 | 2017 | 10.1111/ele.12821 |

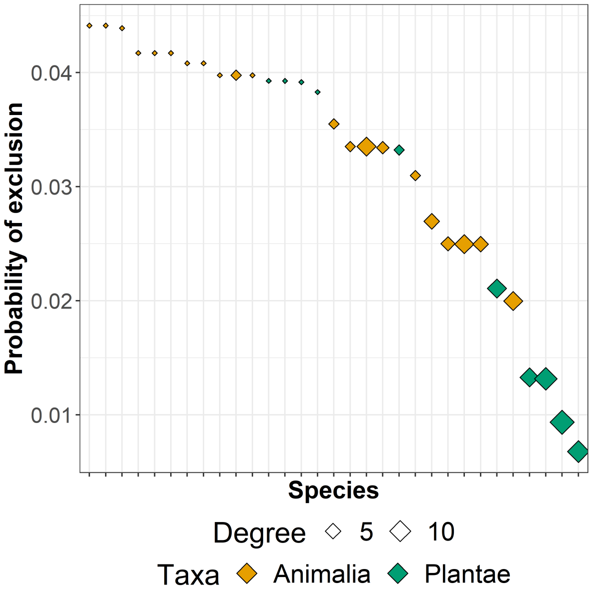


Fig. S16: Probability of exclusion, $P_i^E(\mathbf{A})$ [Eq. (6)], for a mutualistic network in Bartomeus *et al.* (2008) with $S = 31$ species (21 insect and 10 plant nodes, respectively), extracted from the Mangal database (Poisot *et al.*, 2016). To create the interaction matrix, we followed the mean field approximation for intra-guild competition and the two inter-guild parameterizations in (Saavedra *et al.*, 2016) for systems with no interspecific competition, namely: $\rho = 0.00$, $\delta = 0.25$ and $\gamma_0 = 0.50\,\langle\gamma_{ij}\rangle$ (see further details on the parameters in the referred paper). The color and the size of markers show the node's taxa and degree, respectively. Note that for this particular network the authors sampled all plants with an equal sampling effort, hence, plant abundance is not confounded with a larger sampling effort, which may lead to detecting more links, and inflating the degree of abundant plants.

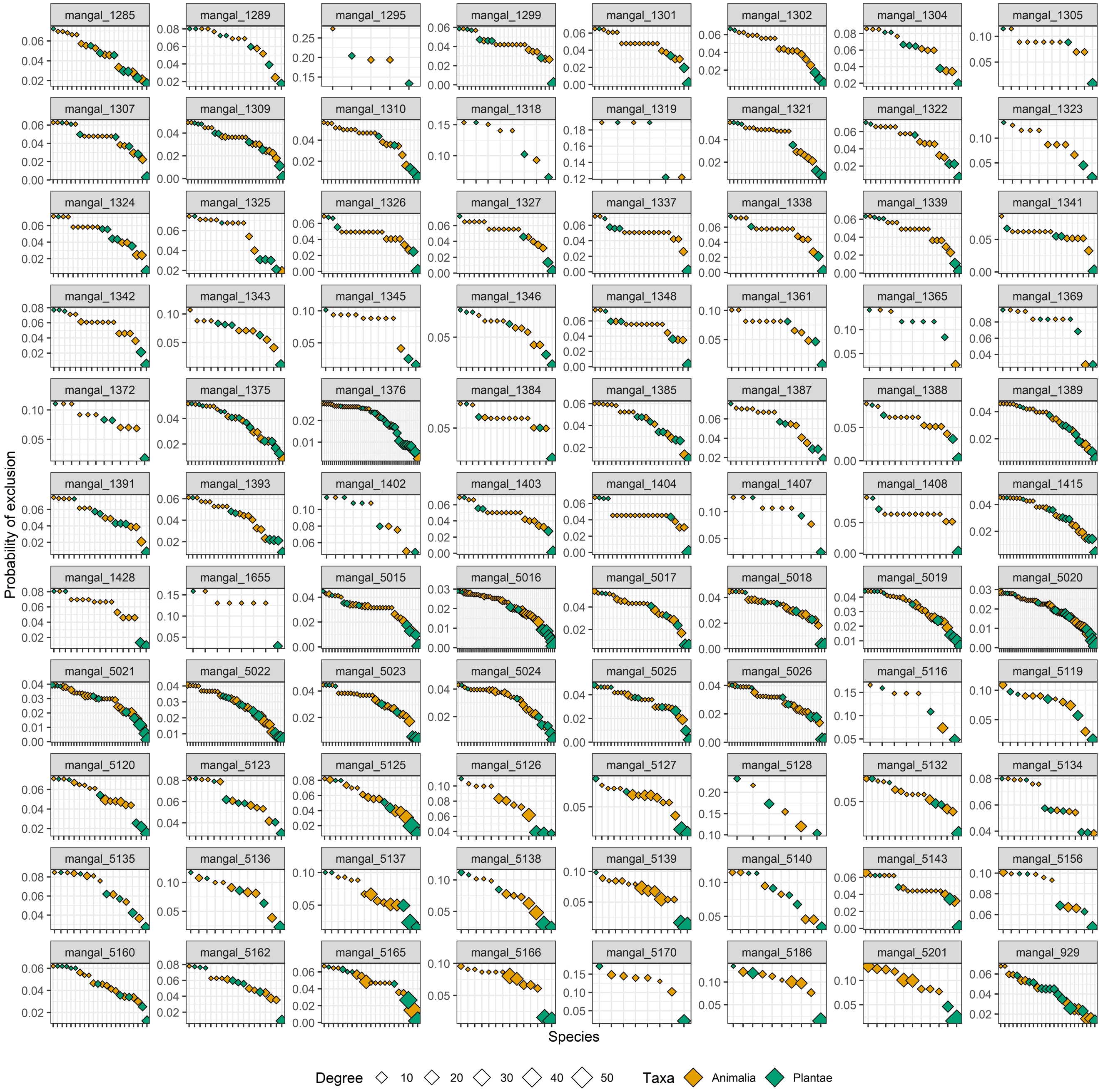


Fig. S17: Probability of exclusion, $P_i^E(\mathbf{A})$ [Eq. (6)], for a sample of 88 bipartite networks, extracted from Mangal (Poisot *et al.*, 2016). To create the corresponding interaction matrices, we used the mean-field approach in (Saavedra *et al.*, 2016) for systems with no interspecific competition, namely: $\rho = 0.00$, $\delta = 0.25$ and $\gamma_0 = 0.50\,\langle\gamma_{ij}\rangle$ (see further details on the parameters in the referred paper). In each network (panel), the color and the size of markers show node's taxa and degree, respectively.

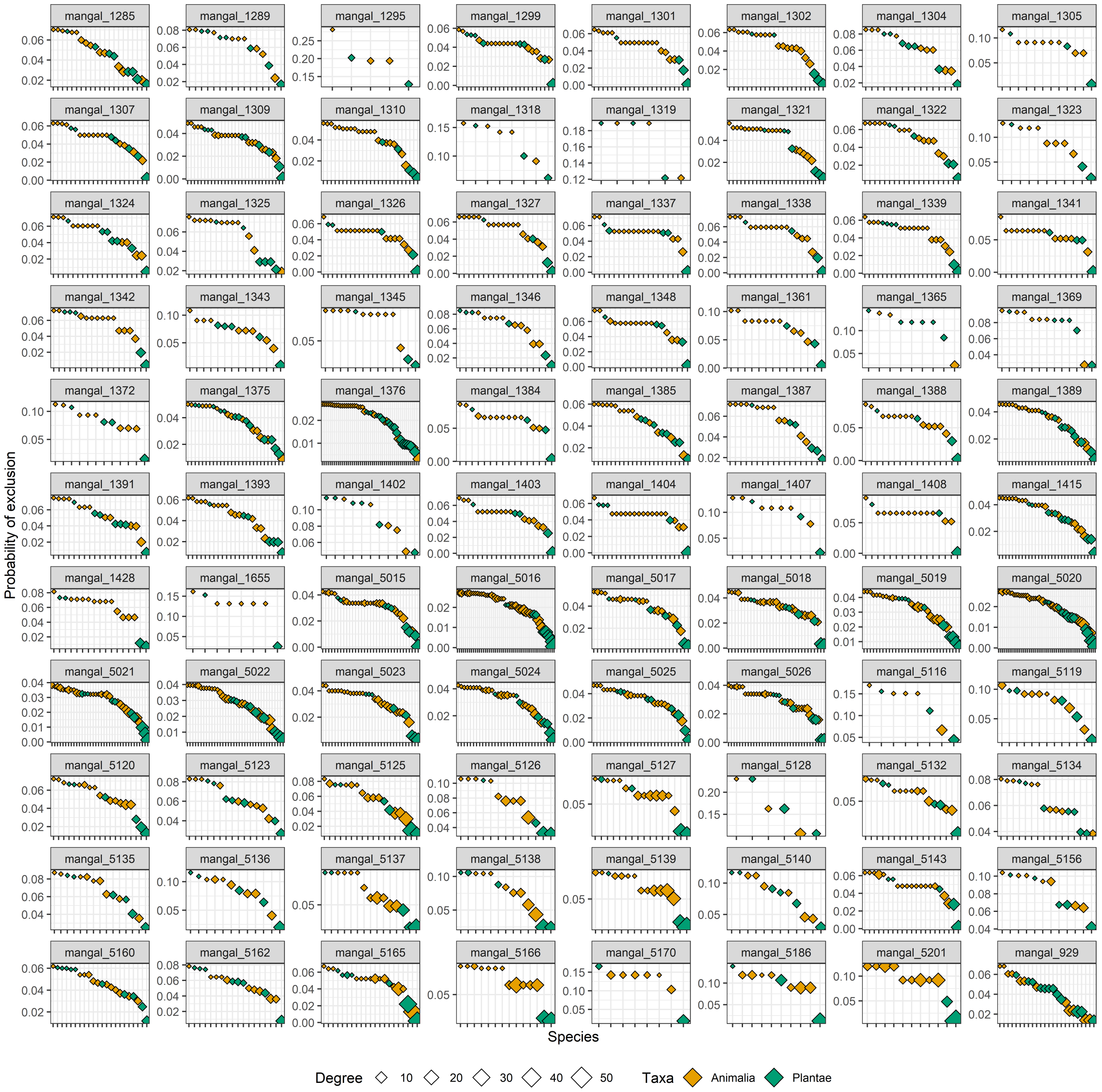


Fig. S18: Probability of exclusion, $P_i^E(\mathbf{A})$ [Eq. (6)], for a sample of 88 bipartite networks, extracted from Mangal (Poisot *et al.*, 2016). To create the corresponding interaction matrices, we used the mean-field approach in (Saavedra *et al.*, 2016) for systems with no mutualistic trade-off, namely: $\rho = 0.01$, $\delta = 0.00$ and $\gamma_0 = 0.50 \langle \gamma_{ij} \rangle$ (see further details on the parameters in the referred paper). In each network (panel), the color and the size of markers show node's taxa and degree, respectively.
